# Cross-species Standardised Cortico-Subcortical Tractography

**DOI:** 10.1101/2025.04.29.651254

**Authors:** Stephania Assimopoulos, Shaun Warrington, Davide Folloni, Katherine Bryant, Ali-Reza Mohammadi-Nejad, Wei Tang, Saad Jbabdi, Sarah Heilbronner, Rogier B Mars, Stamatios N Sotiropoulos

## Abstract

Despite their importance for brain function, cortico-subcortical white matter tracts are underrepresented in diffusion MRI tractography studies. Their non-invasive mapping is more challenging and less explored compared to other major cortico-cortical bundles. We introduce a set of standardised tractography protocols for delineating tracts between the cortex and various deep subcortical structures, including the caudate, putamen, amygdala, thalamus and hippocampus. To enable comparative studies, our protocols are designed for both human and macaque brains. We demonstrate how tractography reconstructions follow topographical principles obtained from tracers in the macaque and how these translate to humans. We show that the proposed protocols are robust against data quality and preserve aspects of individual variability stemming from family structure in humans. Lastly, we demonstrate the value of these species-matched protocols in mapping homologous grey matter regions in humans and macaques, both in cortex and subcortex.

## 1 Introduction

Function-specific brain activity involves the integration of information from multiple remote brain regions. This integration is enabled by white matter (WM) bundles interconnecting different brain regions (*1, 2, 3, 4*). Of particular interest and importance are bundles connecting cortical areas with deep brain structures. Subcortical structures have important roles in affective, cognitive, motor and social functions (*5,6,7*), which emerge through interactions with cortical areas that such connections enable (*8, 9, 10*). Hence, the variability of these connections between individuals has been linked to differences in behavioural traits (*11, 12, 13*). Furthermore, their disruption has been associated with abnormal function and pathology in neurodegenerative and mental health disorders (*14, 15, 16, 17*). In the clinic, individual variability in cortico-subcortical connectivity has been used to assist presurgical planning and predict personalised targets for efficacious interventions (*18*).

Chemical tracer studies in the non-human primate (NHP) brain, have provided, and continue to provide, invaluable insights into cortico-subcortical connections, and neuroanatomy in general (*19*). Examples include tracing of cortico-striatal connections (*20, 21, 8, 22*), amygdalofugal connections (*23*) and thalamo-cortical connections (*24,20*). Comparative neuroanatomy studies can subsequently explore and translate principles of white matter organisation learnt from NHPs to humans (*25, 22, 26*)). Brain imaging and, in particular, diffusion magnetic resonance imaging (dMRI) tractography (*27*) is a crucial component in these comparative studies and beyond (*28, 29, 30*), allowing non-invasive mapping of these connections in the living human.

Towards this direction, recent dMRI-based frameworks have been developed to map respective white matter bundles (“tracts”) across NHPs and humans (*31, 32, 33, 34, 29*). These rely on standardised dMRI tractography protocols, comprising of functionally-driven, rather than geometric, definitions, enabling automated and generalisable mapping of homologous major white matter tracts across species (*31*). These developments have allowed cross-species neuroanatomy studies (*35, 36*) and mapping of connections across humans to study links with brain development, function and dysfunction (*37, 38*). A current limitation of these imaging-based approaches is that they have mainly focused so far on cortico-cortical bundles (with the exception of cortico-thalamic connections).

Tractography protocols for white matter bundles that reach deeper subcortical regions, for instance the striatum or the amygdala, are more difficult to standardise. The relative size and proximity of these bundles, and the white matter complexities and bottlenecks they go through, can make their mapping through dMRI particularly challenging. As a consequence, considerably fewer studies have proposed solutions for their reproducible reconstruction, both within and across primate species, compared to more major cortico-cortical bundles (*39, 40, 31, 41*). Some existing studies have focused on cortico-striatal bundles (*42,43*), uncinate and amygdalofugal fasciculi (*26*), parts of the extreme capsule (*44*) and the anterior limb of the internal capsule (*25,22*). However, these either utilise labour-intensive single-subject protocols (*22,26*), are not designed to be generalisable across species (*42, 43*), or are based mostly on geometrically-driven parcellations that do not necessarily preserve topographical principles of connections (*40*). We propose an approach that addresses these challenges and is automated, standardised, generalisable across two species and includes a larger set of cortico-subcortical bundles than considered before, yielding tractography reconstructions that are driven by neuroanatomical constraints.

Specifically, we build upon our previous work on FSL-XTRACT (*35, 31, 29*) to propose standardised protocols and an end-to-end framework for automated cortico-subcortical tractography in the macaque and human brain, considering connections between the cortex and the caudate, putamen and amygdala. To this end, we use prior anatomical knowledge from NHP tracers to define new generalisable protocols, including the amygdalofugal tract, the Muratoff bundle and the striatal bundle (external capsule) with its frontal, sensorimotor, temporal and parietal parts, augmenting our previous protocols for hippocampal and thalamic tracts (*31*). Due to their close proximity, we also develop new protocols for the respective extreme capsule parts (frontal, temporal, parietal) and revise previously released protocols (*31*) for the uncinate fasciculus, anterior commissure and fornix.

We demonstrate the mapping of the respective bundles in the human and macaque brain and show that tractography reconstructions follow topographical principles obtained from tracers. We show that the proposed definitions are robust against dMRI data quality and preserve aspects of individual variability stemming from family structure in humans, as reflected by higher similarity of reconstructed tracts in the brains of monozygotic twins compared to non-twin siblings and unrelated subjects. We subsequently demonstrate how these tractography reconstructions can improve the identification of homologous grey matter (GM) regions across species, both in cortex and subcortex, on the basis of similarity of grey matter areal connection patterns to the set of proposed white matter bundles (*1, 35*).

## 2 Results

Using prior anatomical knowledge from tracer studies in the macaque, we developed new tractography protocols for the macaque brain and subsequently translated them to the human brain. We considered 23 tracts in total (11 bilateral, 1 commissural), which included tracts connecting the cortex to the amygdala, caudate and putamen. Specifically, we developed protocols for the amygdalofugal (𝐴𝑀𝐹) pathway, the sensorimotor, frontal, temporal and parietal parts of the striatal bundle/external capsule (𝑆𝑡𝐵_𝑚_, 𝑆𝑡𝐵_𝑓_, 𝑆𝑡𝐵_𝑡_, 𝑆𝑡𝐵_𝑝_), and the Muratoff bundle (𝑀𝐵). Due to their proximity, we also developed protocols for the frontal, temporal and parietal parts of the extreme capsule (𝐸𝑚𝐶_𝑓_, 𝐸𝑚𝐶_𝑡_, 𝐸𝑚𝐶_𝑝_) (neighbouring to the corresponding external capsule parts), and revised previous protocols for the uncinate fasciculus (𝑈𝐹) (neighbouring to the 𝐴𝑀𝐹), the fornix (𝐹𝑋) (output tract of the hippocampus next to the amygdala), as well as the anterior commissure (𝐴𝐶) (Table 1 and Supplementary Table S1).

**Table 1:** New and Revised Subcortical Protocols. The developed subcortical tractography protocols for the macaque and human brain. Protocols for Anterior Commissure, Fornix and Uncinate Fasciculus were revised from (31).

| Tract Name | Abbreviation |
| --- | --- |
| Amygdalofugal Tract | $AMF$ |
| Anterior Commissure | $AC$ |
| Extreme Capsule (frontal) | $EmC_f$ |
| Extreme Capsule (temporal) | $EmC_t$ |
| Extreme Capsule (parietal) | $EmC_p$ |
| Fornix | $FX$ |
| Muratoff Bundle | $MB$ |
| Striatal Bundle (sensorimotor) | $StB_m$ |
| Striatal Bundle (frontal) | $StB_f$ |
| Striatal Bundle (temporal) | $StB_t$ |
| Striatal Bundle (parietal) | $StB_p$ |
| Uncinate Fasciculus | $UF$ |

We used the XTRACT approach (*31*) to define tractography protocols, governed by two principles: (i) protocols are comprised of seed/stop/target/exclusion regions of interest (ROIs) defined in template space, so that they are standardised and generalisable (compared to subject-specific protocols), (ii) ROIs are coarse enough and defined equivalently between macaques and humans to enable the tracking of corresponding bundles across species. Full tractography protocols, and modifications to existing protocols, are described in detail in Methods. Protocols were defined in MNI152 template space for human tractography and F99 space (*45, 46*) for macaque tractography. Additionally, we generalised the macaque protocols to the NIMH Macaque Template (NMT v2) (*47*). For clarity, results shown in the main text use the F99 space protocols. Comparison between results in NMT and F99 space can be seen in Supplementary Figure S3.

### 2.1 Subcortical tract reconstruction across species and comparisons with tracers

Using dMRI data from the macaque (𝑁 = 6) and human brain (𝑁 = 50) and the defined protocols, we performed tractography reconstructions for all the tracts of interest. Maximum intensity projections of the resultant group-averaged tract reconstructions for the macaque and human are shown colour-coded in Figure 1 (individual tracts can be seen in Supplementary Figure S1). These reveal overall correspondence in the main bodies of tracts across species, whilst capturing differences in (sub)cortical projections.

**Figure 1:**
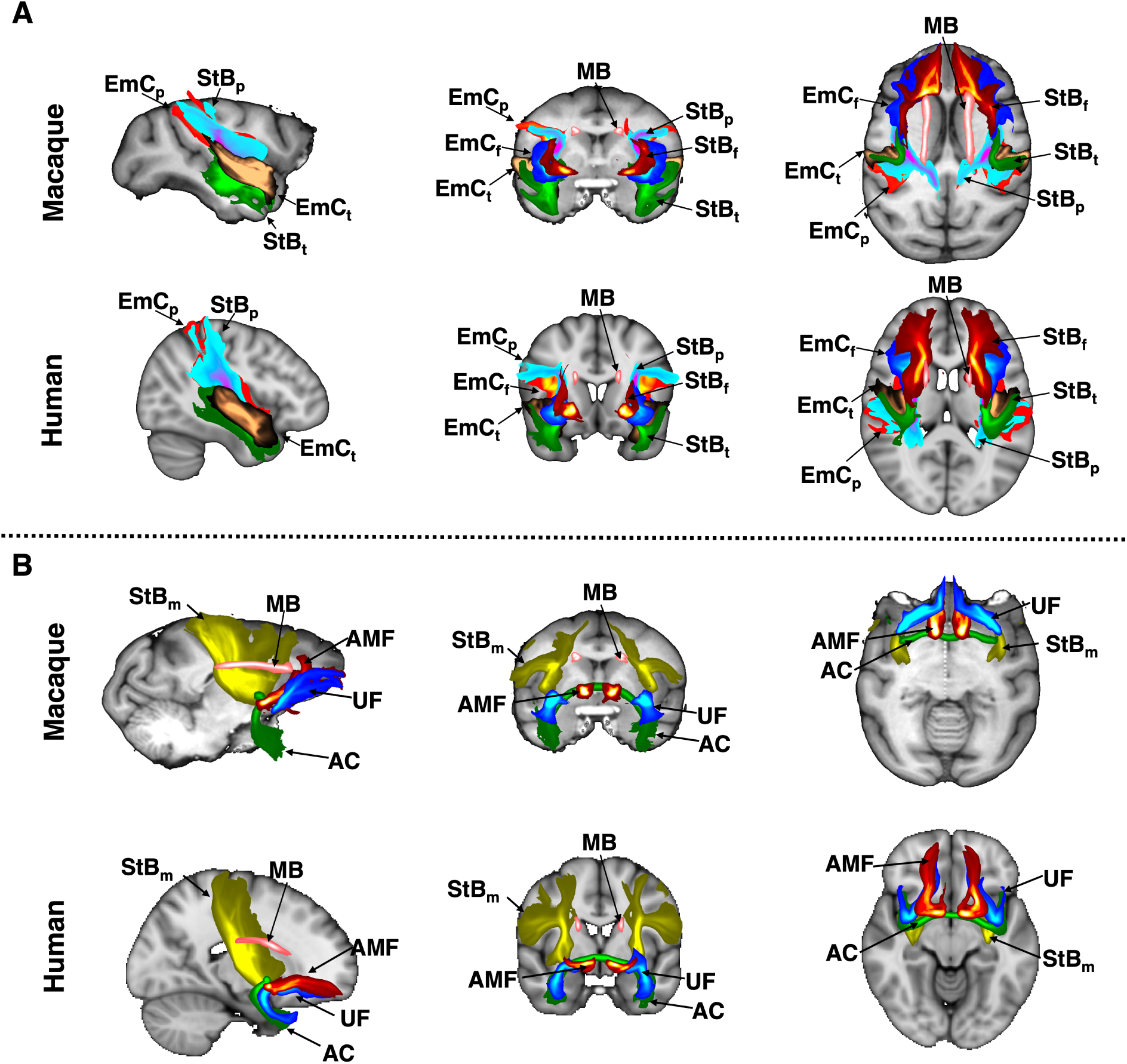
Tractography reconstructions of subcortical bundles in the macaque and human brain using correspondingly defined protocols. Maximum intensity projections (MIPs) in sagittal, coronal and axial views of group-averaged probabilistic path distributions, for all proposed tractography protocols in the macaque (6 animal average) and human (average of 50 subjects from the Human Connectome Project). All MIPs are within a window of 20% of the field of view centred at the displayed slices. (A) Frontal, temporal and parietal parts of the extreme capsule (𝐸𝑚𝐶_𝑓_, 𝐸𝑚𝐶_𝑡_, 𝐸𝑚𝐶_𝑝_); frontal, temporal and parietal parts of the striatal bundle (𝑆𝑡𝐵 _𝑓_, 𝑆𝑡𝐵_𝑡_, 𝑆𝑡𝐵_𝑝_); and the Muratoff bundle (𝑀𝐵). (B) amygdalofugal tract (𝐴𝑀𝐹); anterior commissure (𝐴𝐶); uncinate fasciculus (𝑈𝐹); sensorimotor part of the striatal bundle (𝑆𝑡𝐵_𝑚_); Muratoff bundle (𝑀𝐵). Path distributions were thresholded at 0.1% before averaging.

We explored whether white matter organisation principles known from the tracer literature are captured in these tractography reconstructions (*48,49,20,50,51,10*). For instance, the striatal bundle (𝑆𝑡𝐵)/external capsule is always medial to the extreme capsule and the Muratoff bundle runs along the head of the caudate nucleus. Figure 2A shows the relative positioning for 𝑆𝑡𝐵 _𝑓_, 𝐸𝑚𝐶_𝑓_ and 𝑀𝐵 bundles. Correspondence between tractography results and tract tracing reconstruction in the macaque can be observed, with their relative positions being preserved. This relative position was preserved in the human tractography results as well. Furthermore, the medio-lateral separation is also observed in the other parts of 𝑆𝑡𝐵 and 𝐸𝑚𝐶 (i.e. parietal, temporal), as shown in Supplementary Figure S2. Similarly for the amygdalofugal (𝐴𝑀𝐹) bundle (Figure 2B), this runs through the anterior commissure and ventral pallidum, as well as, in its lateral part, over the uncinate fasciculus (𝑈𝐹). We see agreement with respect to these relative positions in both the macaque and human.

**Figure 2:**
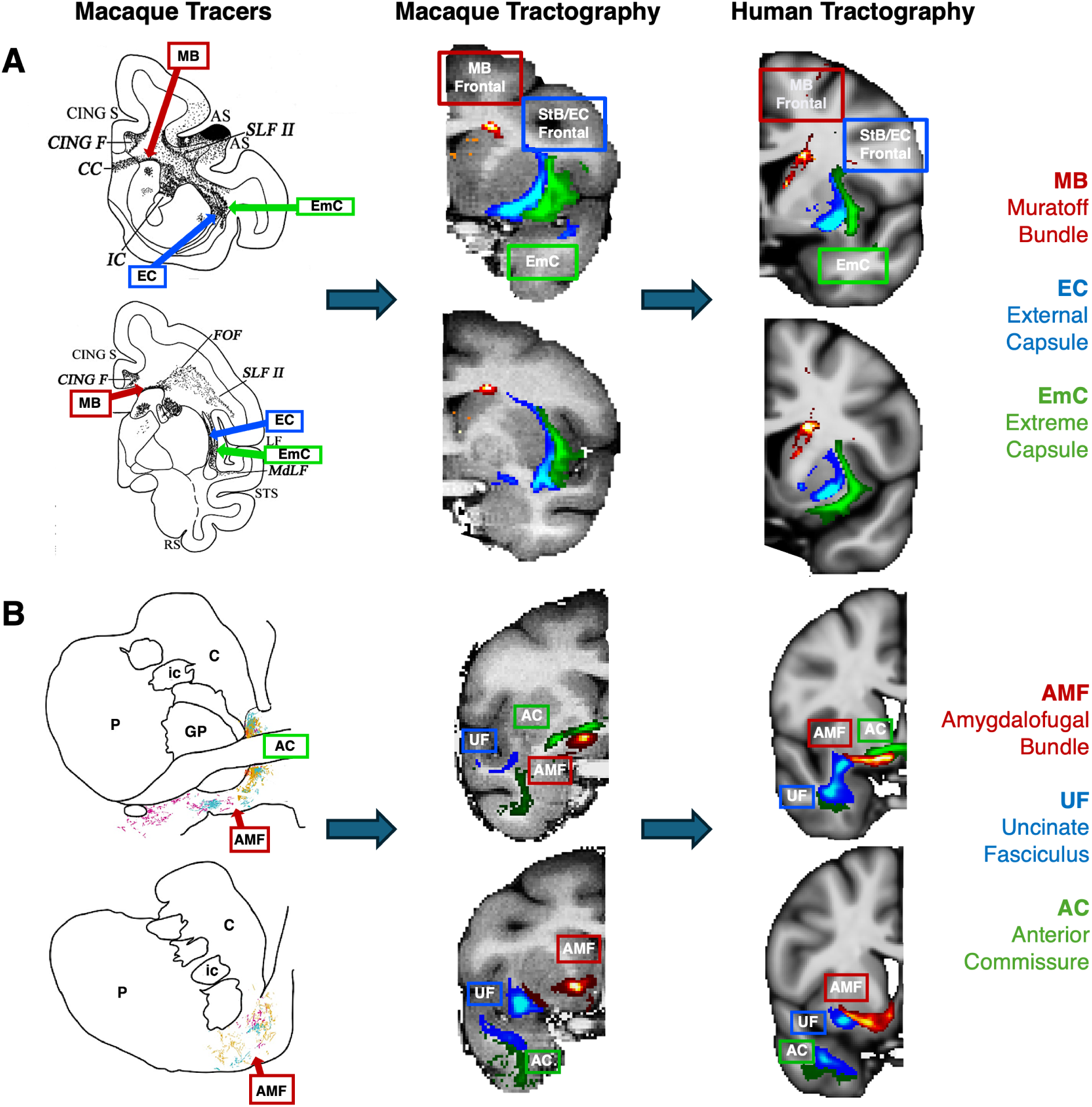
**Tractography mirrors tracer patterns in the macaque brain, with similar patterns in the human**. The proposed protocols were first developed in the macaque guided by tracer literature, and then transferred over to the human. Relative positioning of dMRI-reconstructed tracts was subsequently explored against the ones suggested by tracers, with good agreement in both species. **(A)** The dorsal-medial/ventral-lateral separation between the extreme an external capsule (here the frontal parts 𝐸𝑚𝐶_𝑓_ and 𝑆𝑡𝐵 _𝑓_ shown) is present in macaque tractography, as suggested in the tracer literature. The Muratoff bundle runs along the head of the caudate nucleus. These relative positions are also preserved in the human tractography results. Tracer image modified from (52) with permission. **(B)** Similarly for the amygdalofugal bundle (𝐴𝑀𝐹), which runs under the Anterior Commissure (𝐴𝐶) and over the Uncinate Fasciculus (𝑈𝐹), we see agreement with tracer studies with respect to its location in both the macaque and human tractography (23, 26, 53). Tracer image adapted from (23) with permission. In all examples group-average tractography results are shown.

In addition to the main white matter core of the reconstructed bundles, we also explored agreement of the relative connectivity patterns within the striatum between tracers and tractography. Cortical injections of anterograde tracers from different parts of the macaque brain reveal a dorsolateral to ventromedial organisation in the putamen, from parietal to temporal projections (Figure 3). Using the path distribution of the tractography reconstructed 𝑆𝑡𝐵 parts within the putamen, we could obtain a similar pattern in the macaque brain. This also resembled the pattern found in the human brain, as shown in both coronal and axial views.

**Figure 3:**
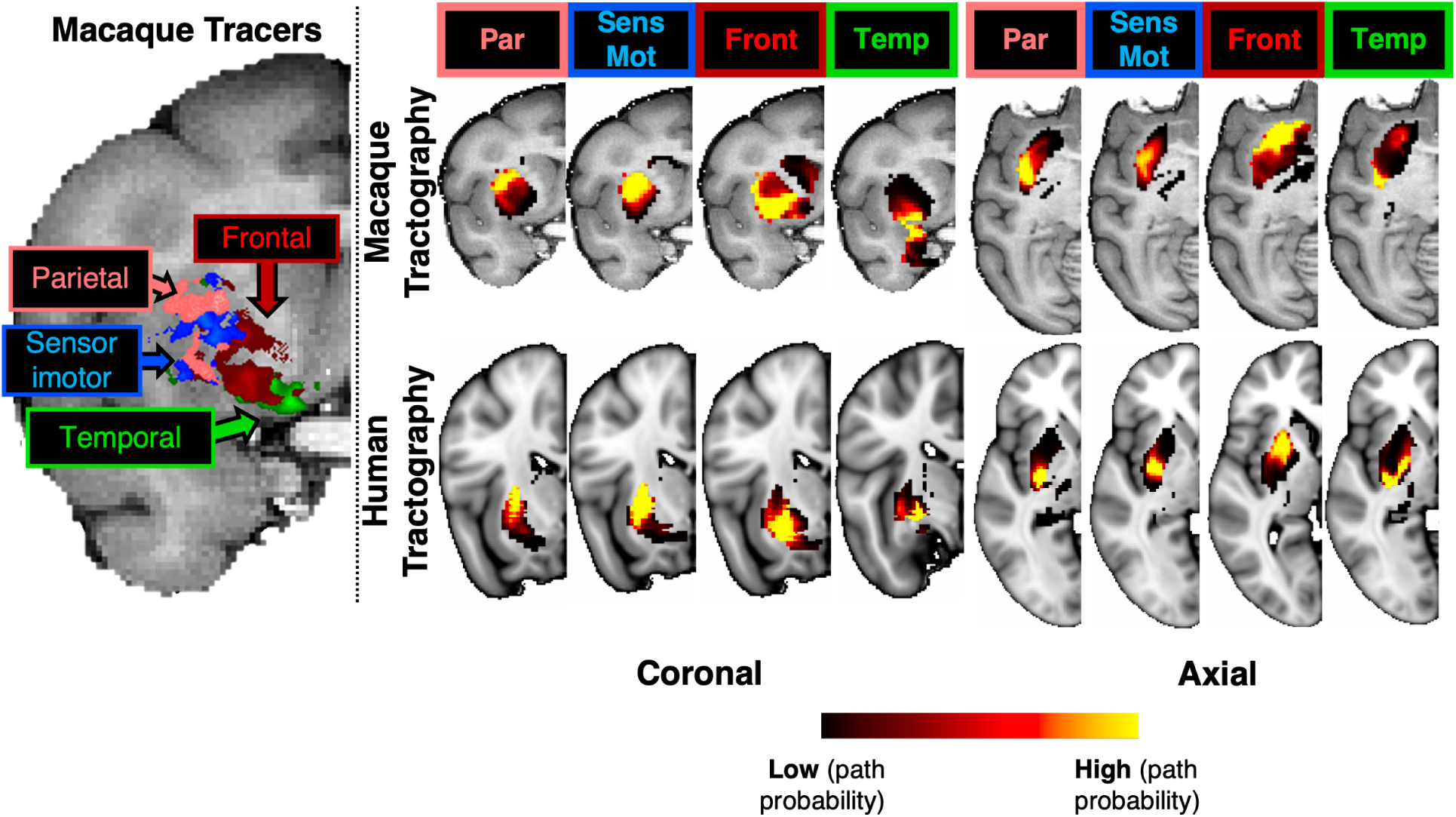
**Tractography-derived connectivity patterns in the putamen resemble (for both macaque and human) termination sites identified by tracers after injections at different cortical areas (frontal, sensorimotor, parietal, temporal) in the macaque. Left**: Using macaque tracer data from 78 injections in various parts of the cortex, tracer termination sites in the putamen suggested a pattern based on the distinct cortical origin of the tracer injection sites; moving from the dorsolateral to the ventromedial putamen. Right: The path distributions of the different parts of the striatal bundle (𝑆𝑡𝐵 _𝑓_, 𝑆𝑡𝐵_𝑚_, 𝑆𝑡𝐵_𝑝_, 𝑆𝑡𝐵_𝑡_) within the putamen reveal a similar pattern of connectivity to different parts of the cortex, both for macaque (top) and the human (bottom). Coronal and axial views of group-average results are shown for tractography. Cortical areas (Front: frontal cortex, Par: parietal cortex, Temp: temporal cortex, SensMot: Sensorimotor cortex) were obtained from the CHARM1 parcellation (54) in the macaque brain (for both tracers and tractography) and from the Harvard parcellation in the human (55, 56, 49).

### 2.2 Generalisability across data and individuals

We subsequently explored generalisability and robustness of the tractography protocols against NHP template spaces and dMRI data quality. Supplementary Figure S3 shows tract reconstructions in the macaque brain when using the F99 (*45*) vs the NMT (*47*) templates, similar tractography reconstructions for protocols defined in either of the two templates.

To explore performance against data quality, we compared tractography reconstruction in very high-quality high-resolution data from the Human Connectome Project (HCP) (*57, 28*), to tractography in more standard quality data from the UK Biobank dataset (*58*). Supplementary Figure S4 demonstrates the ability to reconstruct all tracts across a range of data qualities, with good correspondence of the main bodies of the tracts in both datasets. We quantified this agreement by calculating the mean Pearson’s correlation across the set of new and revised tracts for each unique pair of subjects across and within each of the HCP and UK Biobank (UKB) datasets (Figure 4A). For reference, we performed similar correlations for the original set of XTRACT tracts (*31*) (see Supplementary Table S1 for a list of Original vs New+Revised tracts). Higher correlation was observed within each dataset, but also a sufficiently high correlation between the two datasets. We found similar patterns across datasets both for the original and the new tracts, showcasing that the new protocols behave similarly to the widely-used original XTRACT protocols, across data qualities.

**Figure 4:**
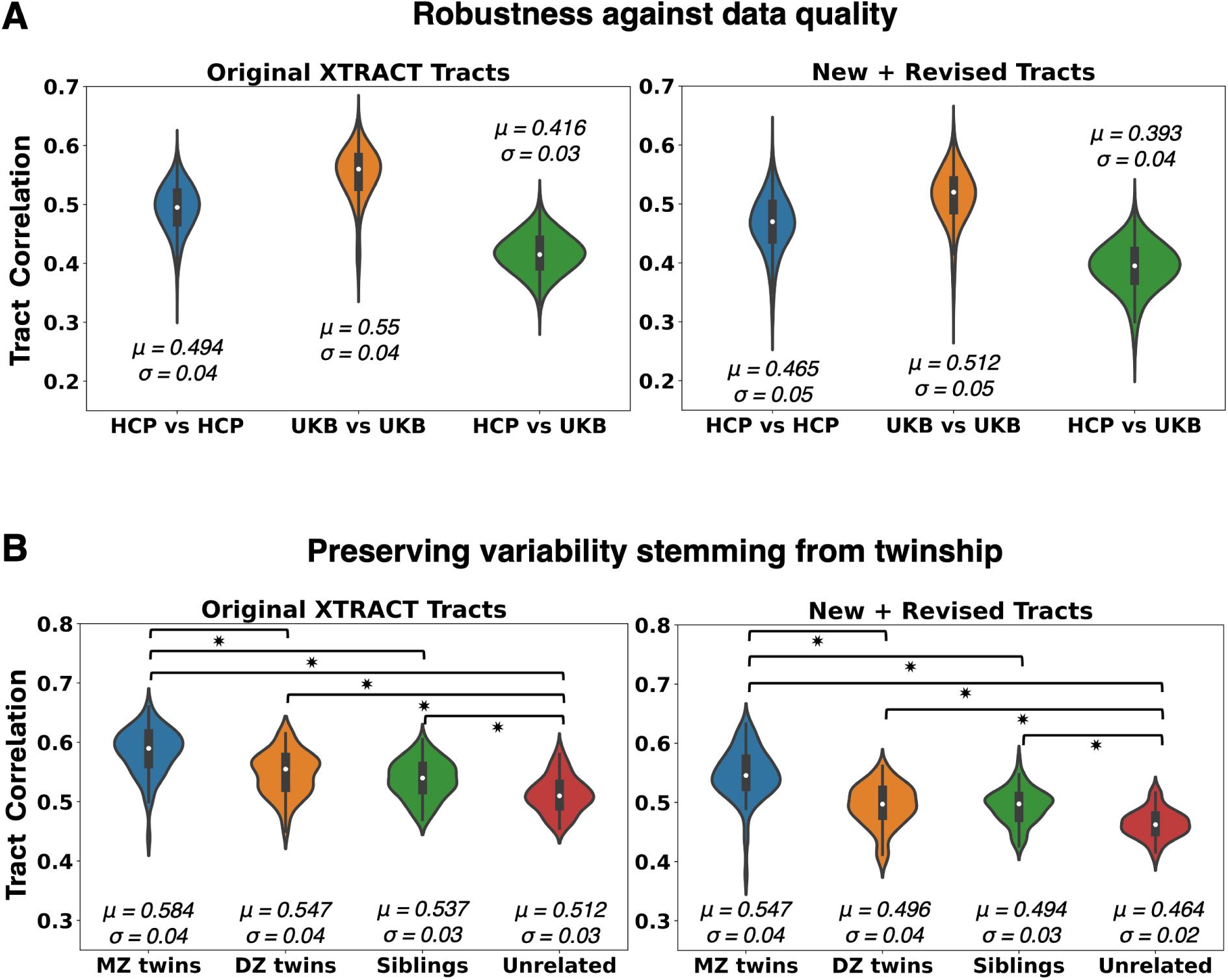
Generalisability of proposed tractography reconstructions across data quality and individuals. Results for the new subcortical tracts (right column) are shown against reference corresponding results for the original set of XTRACT tracts (left column), which have been widely used (31). (**A)** Tract similarity within and between two in-vivo human cohorts, spanning a wide range of dMRI data quality (HCP: high resolution, long scan time, bespoke setup, UK Biobank (UKB): standard resolution, short scan time, clinical scanner). Violin plots of the average across tracts pairwise Pearson’s correlations, between 1,225 unique subject pairs within and across the two cohorts, are shown. Correlations are performed on normalised tract density maps with a threshold of 0.5%. Reported 𝜇 is the mean of the correlations across tracts and subject pairs and 𝜎 is the standard deviation. (**B)** Tract similarity in twins, non-twin siblings and unrelated subjects. Violin plots of the average across tracts pairwise Pearson’s correlations between 72 monozygotic (MZ) twin pairs, 72 dizygotic (DZ) twin pairs, 72 non-twin sibling pairs, and 72 unrelated subject pairs from the Human Connectome Project. Heritable traits are more similar in MZ twins, equally similar in DZ twins and non-twin siblings and more than in unrelated subjects. Asterisk indicates significant pairwise comparisons between groups, as indicated by the brackets.

The mean agreement between HCP and UKB reconstructions was lower compared to within-dataset agreements. The two cohorts correspond to different age ranges, with HCP having younger adults than the UKB, which could be contributing to these differences. In addition, this was due to occasionally reconstructing a sparser path distribution in the low-resolution data, particularly for some of the new tracts, as both their relative size and their proximity make them more challenging. This highlights the potential importance of having high-resolution data in tracking white matter bundles in densely-packed areas of higher complexity. It is interesting to note, however, that similar tracts were less/more reproducible between subjects across data qualities. For the lower quality data from the UKB, the tracts with lowest agreement across subjects (4) were the anterior commissure (𝐴𝐶) and the temporal part of the Extreme Capsule (𝐸𝑚𝐶_𝑡_), while the highest correlations were for the Muratoff Bundle (𝑀𝐵) and the temporal part of the Striatal Bundle (𝑆𝑡𝐵_𝑡_). For the higher quality HCP data, the temporal part of the Extreme Capsule (𝐸𝑚𝐶_𝑡_) and the Muratoff Bundle (𝑀𝐵) were also the tracts with the lowest/highest correlations across subjects, respectively. Hence, certain tract reconstructions were consistently more variable than others across subjects, which may hint to also being more challenging to reconstruct. Taken together, despite differences, our results suggest all tracts could be reconstructed across both data qualities in a generalisable manner (Supplementary Figure S4).

We subsequently explored whether the proposed protocols preserve aspects of individual variability. We used the family structure in the HCP data to explore whether tract reconstructions from monozygotic twin pairs are more similar compared to tracts obtained from other pairs of siblings or unrelated subjects. As shown in Figure 4B, we found a decrease in pairwise tract similarity going from monozygotic twins, to dizygotic twins and non-twin siblings, and to pairs of unrelated subjects. For reference, we performed the same analysis for the original XTRACT tracts and the same pattern persisted for the new (and revised) tracts, in agreement with previous work (*59, 60, 31*). In each analysis, all pairwise differences were significant (Bonferroni corrected 𝑝 ≺ 0.05; following a Mann-Whitney U-Test), with the exception of dizygotic twins compared to non-twin siblings.

Examples of tract reconstructions on individual subjects are shown in Supplementary Figures S6, S7. The figures demonstrate tractography results for subjects corresponding to 10^𝑡ℎ^, 50^𝑡ℎ^ and 90^𝑡ℎ^ percentile of the distribution of tract correlations to the HCP group average. This ranking was also representative of high, medium and low subject motion across the cohort respectively. Results demonstrate that the expected patterns are preserved for all tracts (𝑀𝐵, 𝐴𝑀𝐹, 𝑈𝐹, 𝐸𝑚𝐶, 𝑆𝑡𝐵 (frontal and parietal parts)). Supplementary Figure S6 shows that even the relative medial-lateral organisation of the StB with respect to the EmC is also maintained across the three individual examples, in agreement with the group average pattern.

### 2.3 Identifying homologues in cortex and subcortex using tractography patterns

Based on our previous work (*35*), we used the similarity of areal connectivity patterns with respect to equivalently defined white matter tracts across the two species, to identify homologous grey matter regions between humans and macaques. With the addition of the new subcortical tracts we could perform this task for deep brain structures (subcortical nuclei and hippocampus) with considerably greater granularity than before. Figure 5 demonstrates such identification task for five structures in the left hemisphere (caudate, putamen, thalamus, amygdala, hippocampus) using cortico-cortical and cortico-subcortical tracts (sets of tracts defined in Supplementary Table S1). On the left, the regions in the macaque brain with the lowest divergence (highest similarity) in their connectivity patterns to the connectivity patterns of the corresponding human regions are shown in blue. Using only connectivity pattern similarity, these five structures can be matched almost perfectly across the two species. For instance, human putamen (left hemisphere) has more similar connectivity (lower divergence) to macaque putamen (left hemisphere), human thalamus (left hemisphere) to macaque thalamus (left hemisphere), etc. Since we are mapping structures in the left hemisphere using left hemisphere tracts, we observe a low similarity in the contralateral (right) hemisphere, as expected. On the right of Figure 5, this identification is quantified even further, highlighting the value of considering the new tracts. For every human left-hemisphere region (specified on the vertical axis) the boxplot of divergence of connectivity patterns to each of the five macaque deep brain regions (left-hemisphere) is plotted. The best match corresponds to the boxplot with the lowest values (green) and the dashed blue lines show the medians of these boxplots for each case. For reference, the medians of the divergence values when not considering the new subcortical tracts are shown with the red dashed lines, which are overall more flat (with the exception of the hippocampus which has a connectivity pattern strongly driven by the dorsal subsection of the cingulum bundle (𝐶𝐵𝐷), a cortico-cortical tract). It is evident that considering the new tracts provides enhanced contrast between the subcortical structures connectivity patterns, enabling their correct identification. The improvement is thus not in the best match, but in the specificity of the match.

**Figure 5:**
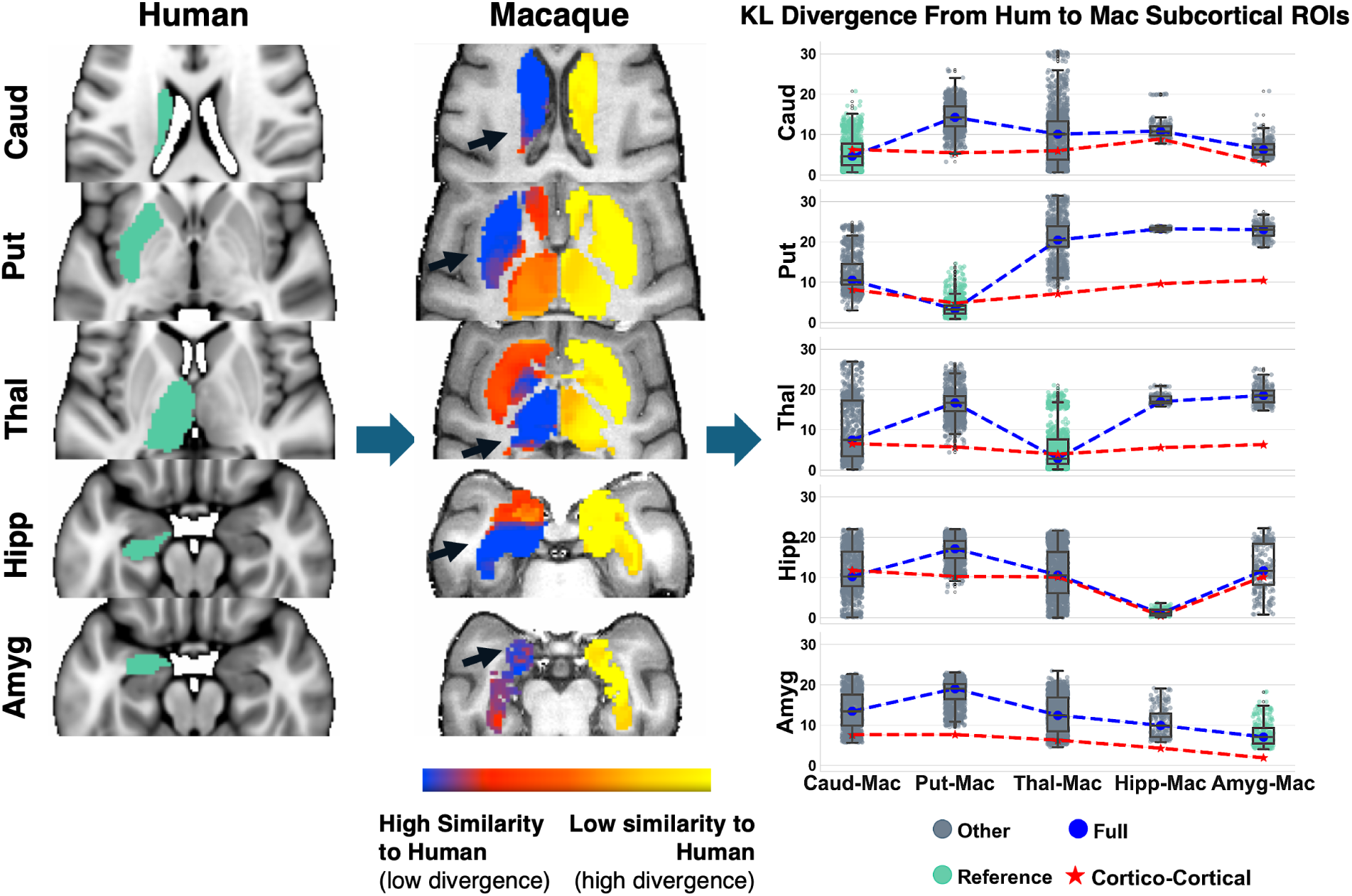
Identifying homologous deep brain structures (subcortical nuclei and hippocampus) across species solely by connectivity pattern similarity, obtained by the new tractography reconstructions. Using the corresponding tracts in humans and macaques, connectivity blueprints can be calculated. These are GMxTracts matrices, with each row providing the pattern of how a GM location is connected to the predefined set of Tracts (35). **Left:** Starting from the average connectivity blueprints of reference human ROIs (Caud: Caudate, Put: Putamen, Thal: Thalamus, Hipp: Hippocampus, Amyg: Amygdala), KullbackLeibler (KL) divergence (or inverse similarity) maps can be computed against the connectivity blueprints of deeper subcortical regions in the macaque (**Middle**). The highest connection pattern similarity corresponds to the homologue macaque region of the corresponding human one. **Right**: Boxplots of KL divergence values between reference human regions and the five macaque ones. Blue dashed line corresponds to median KL divergence values when all white matter tracts are considered (both cortico-cortical and the new subcortical ones). Red dashed line corresponds to median KL divergence when using only cortico-cortical tracts. When cortico-subcortical tracts are included vs not, there is increased specificity/contrast in the cross-species mapping of these deeper structures. The boxplot with the lowest median divergence is shown in green in each case, indicating the best matching regions in the macaque to the human reference (i.e. caudate human reference best matches macaque caudate, putamen human reference best matches macaque putamen, etc).

Having shown increased contrast and specificity in the mapping of deep brain structures, we investigated whether we see a similar effect in the cortex. We selected a set of nearby frontal region pairs to map across the human and the macaque (Figure 6), since a number of the new tracts connect frontal regions to the subcortex. Specifically, we considered the dorsomedial prefrontal cortex (𝑑𝑚𝑃𝐹𝐶), the ventromedial prefrontal cortex (𝑣𝑚𝑃𝐹𝐶), the rostral orbitofrontal cortex (𝑂𝐹𝐶_𝑟_), and the frontal operculum (𝐹𝑂𝑝). These regions were also chosen as they are part of different functional networks (default mode, limbic, and frontoparietal networks), equivalently defined between the macaque and human (*61*).

**Figure 6:**
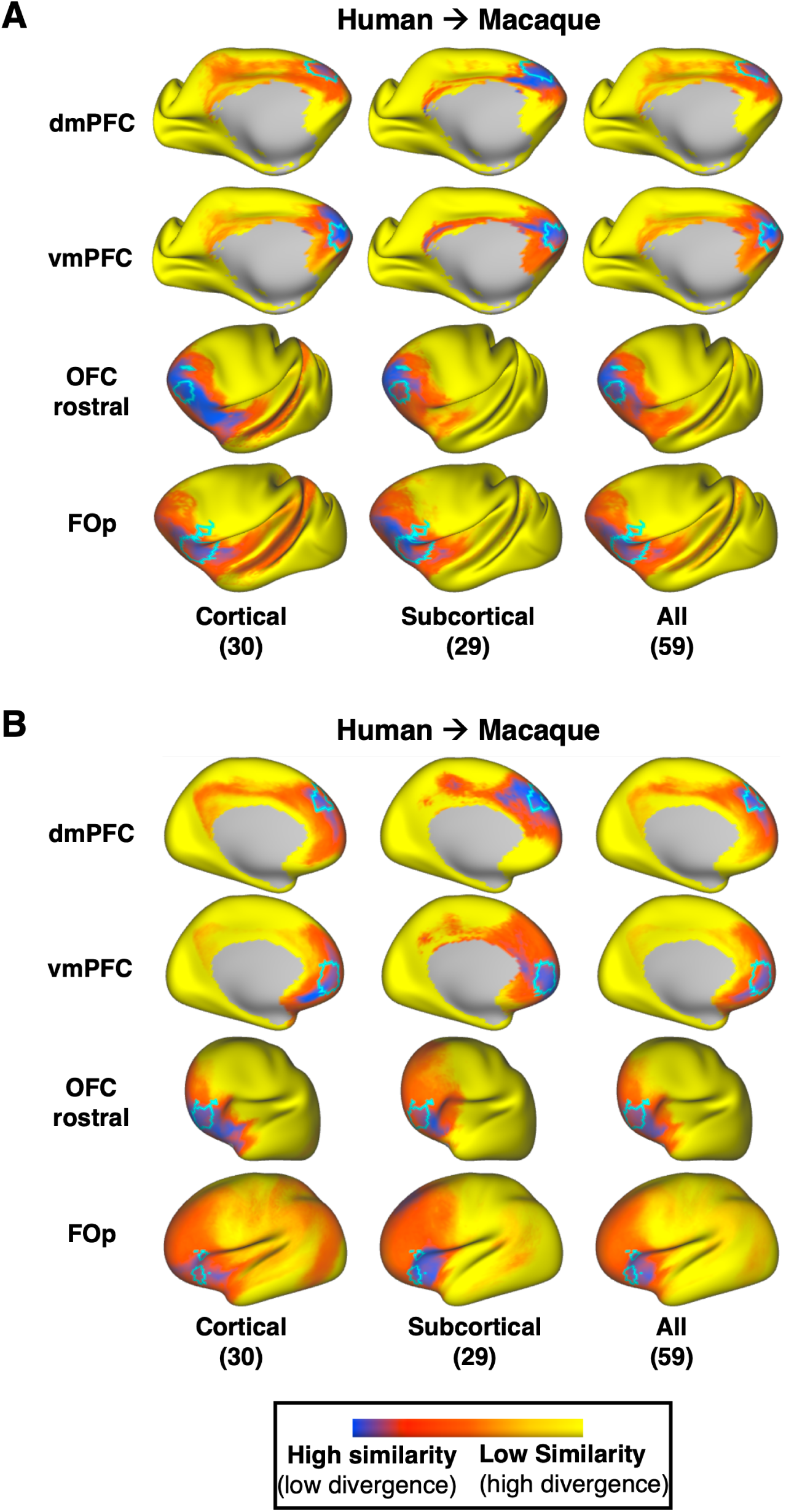
Identifying homologous cortical regions across species solely by connectivity pattern similarity, obtained with and without the new tractography reconstructions. Two pairs of neighbouring frontal regions were chosen (𝑑𝑚𝑃𝐹𝐶: dorsomedial prefrontal cortex & 𝑣𝑚𝑃𝐹𝐶: ventromedial prefrontal cortex, 𝑂𝐹𝐶_𝑟_: rostral orbitofrontal cortex & 𝐹𝑂𝑝: frontal operculum) and their mapping from human to macaque (A) and from macaque to human (B) was explored. For comparison, we overlay in cyan the corresponding homologue regions in each species, as defined in (26). (A) Kullback-Leibler (KL) divergence maps in the macaque for a given human cortical reference region, (one region per row), representing the similarity in connectivity patterns across the macaque cortex to the average pattern of the human reference region. KL divergence maps are calculated using cortico-cortical (1^𝑠𝑡^ column), cortico-subcortical 2^𝑛𝑑^ column) and all tracts (3^𝑟𝑑^ column) to highlight the effect of the cortico-subcortical tractography reconstructions in the prediction. Subcortical tracts provide larger benefits for the prediction of 𝑣𝑚𝑃𝐹𝐶 and 𝑂𝐹𝐶_𝑟_, increasing specificity with respect to the expected borders. (B) Same as in A, but using macaque regions as reference and making predictions on the human cortex. KL divergence maps in the human for a given macaque cortical region, representing the similarity of connectivity pattern across the human cortex to the average pattern of the reference macaque region. Overall, in both species, an increased similarity to the reference regions in the homologue areas and decreased similarity across the rest of the cortex is observed, when cortico-subcortical tracts are considered (2^𝑛𝑑^ or 3^𝑟𝑑^ column).

The prediction from human to macaque is shown in Figure 6A, while the converse prediction from macaque to human is shown in Figure 6B. In each case, we compare the prediction using only cortico-cortical tracts (column 1), using only cortico-subcortical tracts (column 2), and using the full set (column 3) (sets of tracts defined in Supplementary Table S1 - the middle cerebellar peduncle (𝑀𝐶𝑃) was not used in these comparisons). The predicted areas with the highest similarity in connectivity patterns are depicted in blue, while the a priori expected homologue region borders have been outlined in cyan. These results demonstrate benefits when using the subcortical tracts, with mapping of some regions (for instance 𝑣𝑚𝑃𝐹𝐶 and 𝑂𝐹𝐶_𝑟_) being improved more than others. However, in general, we observed an increase in cross-species similarity in the corresponding areas of interest, combined with a decrease in similarity everywhere else in the cortex, when we considered cortico-subcortical tracts (columns 2, 3) compared to when we considered cortico-cortical tracts alone (column 1).

Figure 7 provides a further insight into these mappings, by plotting the connectivity patterns of the human regions against the pattern of its identified best match in the macaque brain. As can be observed, despite the relative proximity of these frontal regions, we have distinct patterns across them. With the exception of 𝑑𝑚𝑃𝐹𝐶, the connectivity patterns of all other regions have major contributions from the frontal striatal bundle, the extreme capsule and the amygdalofugal tract (𝐴𝑀𝐹) and connection patterns to these cortico-subcortical bundles enable better separation of these nearby regions. For instance, 𝑣𝑚𝑃𝐹𝐶 and 𝑑𝑚𝑃𝐹𝐶 have both connections through the cingulum bundle, the corpus callosum and the inferior fronto-occipital fasciculus. However, they connect differently to the striatal bundle, the amygdalofugal tract and the anterior thalamic radiation and the addition of these tracts in the connectivity patterns allow the two regions to be better distinguished. The 𝐹𝑂𝑝 and 𝑂𝐹𝐶_𝑟_ have both relatively strong connection patterns to the uncinate and the inferior fronto-occipital fasciculi, but it is their different pattern of connections to extreme and external capsules and the amygdalofugal tract that enable their better separation.

**Figure 7:**
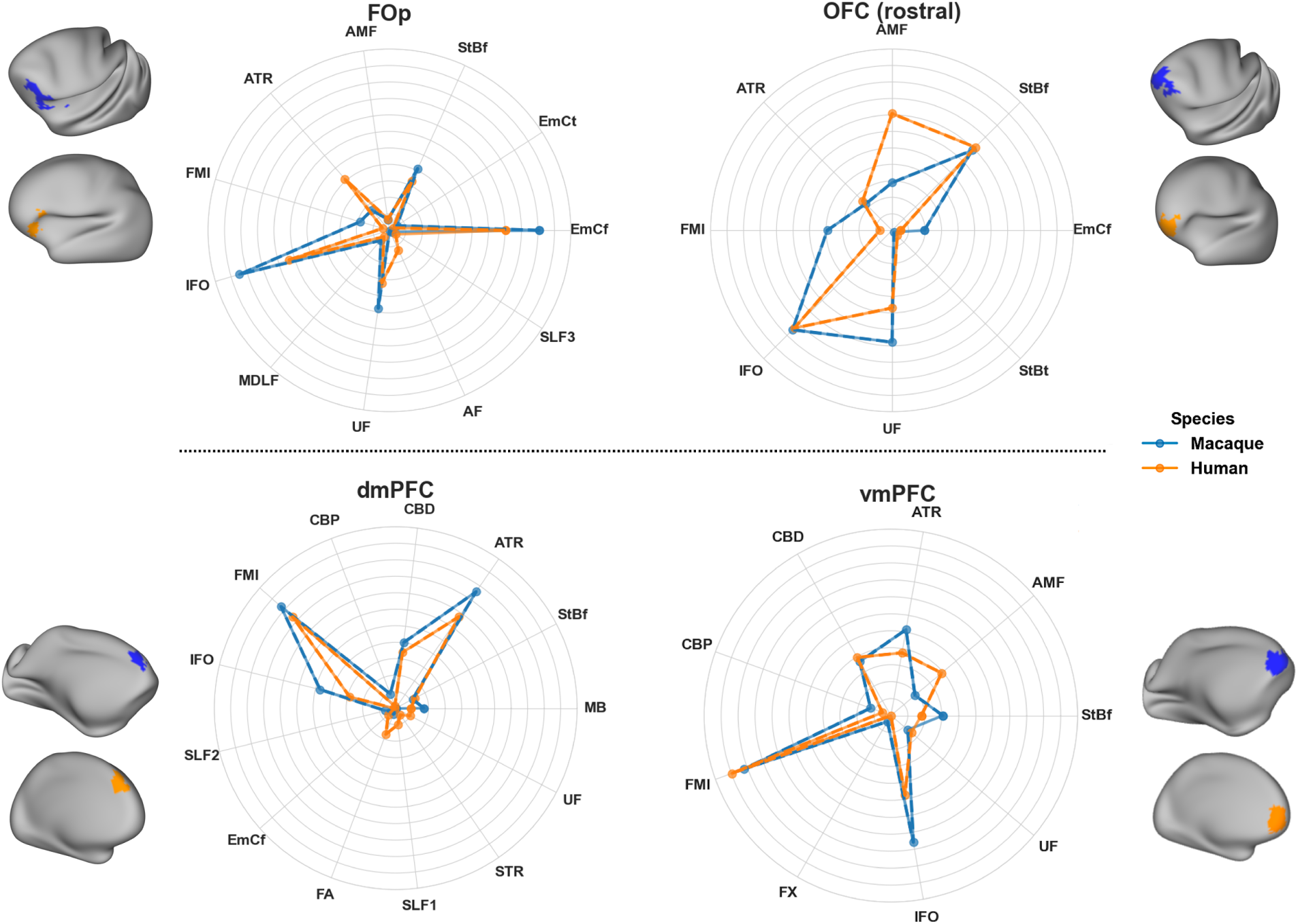
Connectivity patterns for neighbouring frontal region pairs, showing distinct cortico-subcortical tract contributions in macaque and human. Considered regions are the same as in Figure 6A, i.e. 𝐹𝑂𝑝: frontal operculum, 𝑂𝐹𝐶_𝑟_: rostral orbitofrontal cortex, 𝑑𝑚𝑃𝐹𝐶: dorsomedial prefrontal cortex, 𝑣𝑚𝑃𝐹𝐶: ventromedial prefrontal cortex. Reference regions were chosen in the human cortex, shown in orange, and obtained from (26). The best matching region across the whole macaque cortex was identified by the minimum KL divergence in connectivity patterns (thresholded at the 7^𝑡ℎ^ percentile in each case) and is shown in blue. Average connectivity patterns for the reference and best-matching regions are depicted using the polar plots. For each region, similarities in the connectivity patterns between the macaque and human can be observed, with the new cortico-subcortical bundles contributing to these patterns. For instance, 𝐹𝑂𝑝 has a strong connection pattern involving 𝐸 𝑀𝐶_𝑓_ and 𝑈𝐹 and moderately 𝑆𝑡𝐵 _𝑓_ and 𝐴𝐹, while its neighbouring 𝑂𝐹𝐶_𝑟_has a stronger pattern involving 𝑆𝑡𝐵 _𝑓_ and 𝑈𝐹, compared to 𝐸 𝑀𝐶_𝑓_. These differences are preserved across both species.

## 3 Discussion

We introduced standardised dMRI tractography protocols for delineating cortico-subcortical connections between cortex and the amygdala, caudate, putamen and the hippocampus, across humans and macaques. Building upon our previous work (*35, 31*), which already provided protocols for cortico-thalamic radiations, and guided by the chemical tracer literature in the macaque, we devised the new protocols first for the macaque and then extended to humans. We demonstrated that our reconstructed tracts preserve topographical organisation principles, as suggested by tracers (*62,8,63*). As outlined in (*43*), tractography reconstructions can be highly accurate if information about where pathways go, and where they do not go is available. This is the philosophy behind the proposed protocols, which provide this type of constraints across different bundles. At the same time these constraints are relatively coarse so that they are species-generalisable. We found that the proposed approaches yield tractography reconstruction across a range of datasets and respect individual similarities stemming from twinship. We further assessed the efficacy of these protocols in performing connectivity-based identification of homologous cortical and subcortical areas across the two species (*35, 36, 38*).

Mapping white matter tracts that link cortical areas with deep brain structures (subcortical nuclei and hippocampus), as done here, enhances capabilities for studying neuroanatomy in many contexts, from evolution and development, to mental health and neuropathology. As one of the (evolutionarily) older brain structures, the subcortex modulates brain functions including basic emotions, motivation, and movement control, providing a foundation upon which the more complex cognitive abilities of the cortex could develop and evolve (*62,64,8,65,66*). This modulatory function is mediated via white matter bundles (*8, 9*). Consequently, their disruption is linked to abnormal function and pathology, in mental health, neurodegenerative, and neurodevelopmental disorders (*15, 67, 17*). For example, in depression, fronto-thalamic (*68*), cortico-amygdalar (*69, 70*), and corticostriatal (*71*) connectivity changes have been reported, while in schizophrenia there are associated fronto-striatal (*72*) and hippocampal connectivity (*73*) changes. In Parkinson’s there is impairment in fronto-striatal connectivity (*74, 75, 76, 77*), while fronto-thalamic and cingulate connectivity are impaired in Alzheimer’s disease (*75, 78*). Connectivity between the frontal lobe and the amygdala, thalamus, and striatum, as well as cingulum connectivity are impaired in obsessive compulsive disorder (OCD), autism spectrum disorder (ASD), and attention deficit hyperactivity disorder (ADHD) (*79, 69, 14, 80*). Therefore, reconstructing connectivity of these deep brain structures (striatum, thalamus, amygdala, hippocampus) in a standardised manner, as enabled by our proposed tools, allows for further investigation into a wide range of disorders.

In addition, tractography of connections linking to/from deep brain structures has been used or proposed for guiding neuromodulation interventions, for example, deep brain stimulation (DBS) (*81, 82*) or repetitive transcranial magnetic stimulation (rTMS) (*67*). DBS can inherently target subcortical structures and connectivity of subcortical circuits can be used to identify efficacious stimulation targets (*83, 18*). rTMS on the other hand modulates subcortical function indirectly by targetting the structurally connected cortical areas. For example, dmPFC has been targetted to modulate the reward circuitry, in cases of anhedonia, negative symptoms in schizophrenia and major depression disorder (MDD) (*84, 85, 86*), while the vmPFC has been used as a target to modulate the prefrontal-striatal network (part of the limbic system) and regulate emotional arousal/anxiety (*87, 88, 89*). Our results show a good mapping across species of both these cortical regions with specificity in their connectional patterns. Additionally, the motor cortex has been used as target to modulate cortico-striatal connectivity in general anxiety disorder (*90, 91*). We thus anticipate that having a standardised set of tracts linking the striatum, the hippocampus, the amygdala and the thalamus (all potential sites for stimulation) to specific cortical areas can assist the planning of interventions.

Our cross-species approach naturally lends itself to the study of evolutionary diversity. A number of comparative studies have revealed differences and similarities when comparing brain connectivity between humans and non-human primates (*92*), including macaques (*35, 38*) and chimpanzees (*93*). Our work naturally extends these efforts and provides new tools for studying this diversity in deeper structures and subcortical nuclei. The ever increasing availability of comparative MRI data (*34, 94*) allows the definition of similar protocols in more species, such as the gibbon (*33, 95*) or the marmoset monkey, and even across geometrically diverse brains depicting different stages of neurodevelopment (e.g. neonates vs adults) enabling concurrent studies of phylogeny and ontogeny (*38*)).

Our protocols have been developed and tested using FSL-XTRACT, but, in principle, are not specific to FSL. We have not evaluated performance with other tools, but these standard space protocols could be translated into other tractography approaches. As described before, the protocols are recipes with anatomical constraints, including regions to which the corresponding white matter pathways connect and regions they do not, constructed with cross-species generalisability in mind. Caution may be needed, however, if applying such protocols for segmenting whole-brain tractograms, as these can induce more false positives than tractography reconstructions from smaller seed regions and may require stricter exclusions.

Despite the potential demonstrated in this work, our study has limitations. As this is the first endeavour of this scale to map cortico-subcortical connections in a standardised manner and across two species, it is not exhaustive. Tracts linking the cortex to the striatum were prioritised as they are of increased relevance in human development and disease. However, expanding to include more tracts targetting other structures would provide a more holistic view. Our protocols were developed in the adult human brain. Future work will translate them to the infant brain (expanding on previous work (*38*)) to interrogate cortico-subcortical connectivity across development. Tractography validation is a challenge, as is validation for any indirect and non-invasive imaging approach. We explored and demonstrated generalisability of the proposed protocols, both within and across species. We also showed how the imaging-based reconstructions follow topographical organisation principles suggested by tracers.

## 4 Methods and Materials

### 4.1 Tractography protocols

Guided by tract tracing and neuroanatomy literature, we devised tractography protocols for 18 subcortical bundles (nine bilateral - Table 1) using the XTRACT approach (*31*). We also revised protocols for three more bundles (2 bilateral, 1 commissural), compared to their original version (*31*). All protocols followed two principles: i) comprised of seed/stop/target/exclusion regions of interest (ROIs) defined in template space, so that they are standardised and generalisable, ii) ROIs defined equivalently between macaques and humans to enable tracking of corresponding bundles across species. The human protocols were defined in MNI152 space. The macaque protocols were defined in F99 and also in NMT space.

The tracts included the amygdalofugal tract (𝐴𝑀𝐹), the uncinate fasciculus (𝑈𝐹), anterior commissure (𝐴𝐶), sensorimotor, temporal, parietal and frontal parts of the striatal bundle (𝑆𝑡𝐵) / external capsule (𝐸𝐶), Muratoff bundle (𝑀𝐵)/subcallosal fasciculus, as well as the extreme capsule (𝐸𝑚𝐶) parts that run close to the putamen connecting the insula to the frontal, temporal and parietal cortices. All XTRACT tracts (Original, Revised and New) are summarised in Supplementary Table S1.

Detailed protocol definitions are presented below and summarised in Figure 8 (for completeness, the previously published thalamic radiations from (*31*) are presented in Supplementary Materials). Briefly, the 𝐴𝑀𝐹 and 𝑈𝐹 protocols are a standard-space generalisation of the individual subject-level protocols presented in (*26*). For the remaining protocols, we first devised them in the macaque guided by tract tracer literature. Specifically, the approach we took was to first identify anatomical constraints from neuroanatomy literature for each tract of interest independently, derive and test these protocols in the macaque. Thus, each devised protocol included a unique combination of anatomically defined masks (based on literature descriptions of the tracts), delineated in standard macaque space (F99). We then developed corresponding protocols in the human using correspondingly defined landmarks (delineated in standard MNI space). We optimised in an iterative fashion based on two criteria: (1) the protocols generalise well to humans, and (2) when considering groups of bundles, the generated reconstructions follow topographical principles known from tract tracing literature.

**Figure 8:**
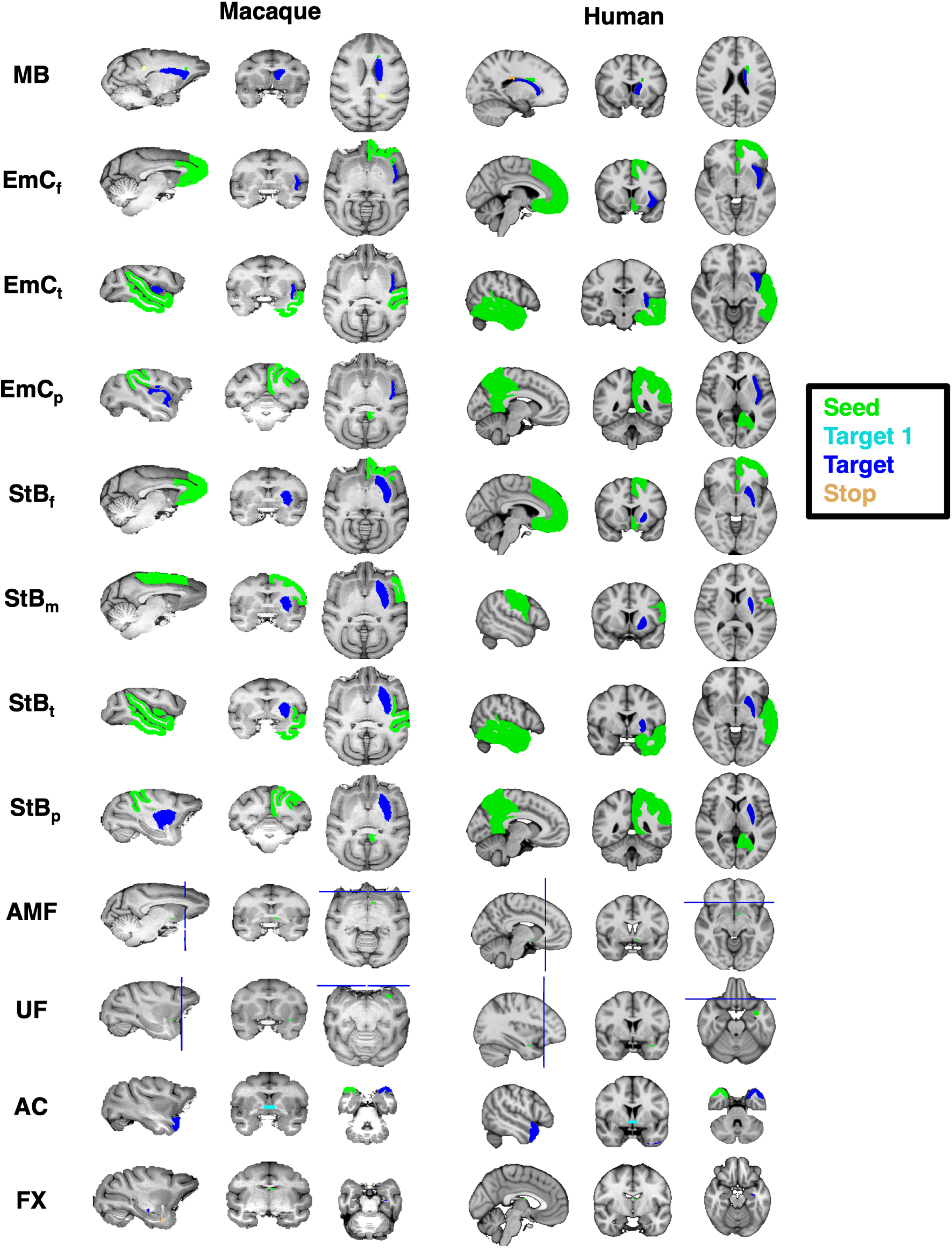
Corresponding tract protocol definitions across species. Protocol definitions for all new (and revised) tracts in the human and macaque. Protocols were first designed in the macaque brain guided by macaque tracer literature, and then transferred over to the human. Colour-coded regions depict the seed, target and stop masks. Exclusion masks are not shown for ease of visualisation.

We modified existing XTRACT protocols to improve their specificity in the subcortex. Specifically, we developed a new uncinate fasciculus (𝑈𝐹) protocol based on the protocol presented in (*26*). We also modified the anterior commissure (𝐴𝐶) protocol to improve temporal lobe projections and slightly enhance projections to the amygdala, and modified the fornix (𝐹𝑋) one to reduce amygdala projections by placing an exclusion in the amygdala.

Given the proximity of the newly defined tracts, we evaluated the new protocols against their ability to capture patterns known from the tracer literature. These included relative positioning of each tract with respect to neighbouring tracts (Figure 2) and topographical organisation of certain bundle terminals within subcortical nuclei (Figure 3).

#### 4.1.1 New protocol definitions

##### Amygdalofugal Pathway (𝐴𝑀𝐹)

We derived generalisable template-space protocols to reconstruct the limbic-cortical ventral amygdalofugal pathway, following the subject-specific protocols in (*26*). The amygdalofugal pathway (𝐴𝑀𝐹) courses between the amygdala and the prefrontal cortex (PFC), running alongside the uncinate fasciculus (𝑈𝐹) medially and finally merging with the 𝑈𝐹 in the posterior orbitofrontal cortex (OFC). As in (*26*), the seed included voxels with high fractional anisotropy in an anterior-posterior direction in the sub-commissural white matter. We used a target covering all brain at the level of caudal genu of the corpus callosum (same target as in the revised 𝑈𝐹 protocol, described further below). Exclusions include an axial plane through the 𝑈𝐹, the internal and external capsules, the corpus callosum, the cingulate, the Sylvian fissure, the anterior commissure, the fornix, and a large coronal exclusion covering all brain dorsal to the corpus callosum and extending inferiorly to the frontal operculum and insula at the level of the middle frontal gyrus.

##### Striatal Bundle (𝑆𝑡𝐵 _𝑓_, 𝑆𝑡𝐵_𝑚_, 𝑆𝑡𝐵_𝑝_, 𝑆𝑡𝐵_𝑡_)

The striatal bundle (𝑆𝑡𝐵), is a bundle system that connects the cortex to the striatum, and joins the external capsule (𝐸𝐶). Although terminations reach both the caudate and the putamen, it primarily terminates in the putamen (*48,49,10*). Here we defined protocols for bundles that connect the putamen with frontal (including anterior cingulate) lobe, sensorimotor cortex, parietal and temporal lobes. For all parts, we used the putamen as the target. For 𝑆𝑡𝐵 _𝑓_, the OFC, PFC, ACC and frontal pole made up the seed. For 𝑆𝑡𝐵_𝑚_, the primary sensorimotor cortex (M1-S1) were used as seeds. For 𝑆𝑡𝐵_𝑡_ and 𝑆𝑡𝐵_𝑝_, the temporal and parietal lobes were respectively used as a seed. The exclusion masks shared many commonalities but also had differences. For all parts, exclusions included a midsagittal plane, the subcortex, except for the putamen, as well as the occipital lobe. For each of the parts, we additionally excluded the seeds for every other 𝑆𝑡𝐵 bundle.

##### Muratoff Bundle (𝑀𝐵)

The subcallosal fasciculus tract (as called in human neuroanatomy) or Muratoff bundle (as called in non-human animal neuroanatomy) (*48, 51*) is a complex system of projection fibres which runs beneath the corpus callosum, above the caudate nucleus at the corner formed by the internal capsule and the corpus callosum (*42*). Although terminations reach both the caudate and the putamen, it primarily terminates in the caudate head (*48, 42, 51*). As its cortical projections are challenging to capture and isolate using tractography, we defined a protocol for the major core of the bundle. We used a seed in the white matter adjacent to the caudate head and a target in the white matter adjacent to the caudate tail. A stop mask was used beyond the target in the white matter above the target. Exclusions included the contralateral hemisphere, the subcor-tex (except for the caudate head), the brainstem, the parietal, occipital, frontal and temporal cortices.

##### Extreme Capsule (𝐸𝑚𝐶_𝑓_, 𝐸𝑚𝐶_𝑝_, 𝐸𝑚𝐶_𝑡_)

The extreme capsule is a major association fascicle that carries association fibres between frontal-temporal and frontal-parietal, as well as these areas and the insula (*49*). It lies between the claustrum and the insula, with the claustrum being considered the boundary between the 𝐸𝑚𝐶 and the external capsule (𝐸𝐶) (*10*). We defined protocols connecting the insula to frontal, parietal and temporal cortices. For all parts we used the insula as the target, while for seeds we used the same seeds as for the corresponding 𝑆𝑡𝐵 parts. Hence, for the 𝐸𝑚𝐶_𝑓_protocol the frontal pole was used as the seed. For 𝐸𝑚𝐶_𝑝_, the parietal lobe was a seed, and for 𝐸𝑚𝐶_𝑡_ the temporal lobe was a seed. Exclusions for all 𝐸𝑚𝐶 parts included the contralateral part of the brain, the subcortex, as well as the occipital lobe, and lateral parts of the somatosensory and motor cortices. In addition, for each subdivision, the exclusion mask also included the seed mask for every other subdivision.

#### 4.1.2 Revisions to previous XTRACT protocols

##### Uncinate Fasciculus (𝑈𝐹)

The 𝑈𝐹 lies at the bottom part of the extreme capsule, curving from the inferior frontal cortex to the anterior temporal cortex. Given the neighbouring bundles that were newly defined, we took a new approach to the 𝑈𝐹 compared to the original XTRACT implementation (*31*), now following the principles of (*26*). Briefly, we used an axial seed in the white matter rostro-laterally to the amygdala in the anterior temporal lobe. A target covered all brain at the level of caudal genu of the corpus callosum. Exclusions included the basal ganglia, a coronal plane posterior to the seed, the corpus callosum, the cingulate, the Sylvian fissure, the anterior commissure, and a large coronal exclusion covering all brain dorsal to the corpus callosum and extending inferiorly to the frontal operculum and insula at the level of the middle frontal gyrus. This implementation provided improved connectivity to the dorsal frontal cortex and aided separability with respect to neighbouring white matter bundles.

##### Anterior Commissure (𝐴𝐶)

Compared to original XTRACT protocol, we entirely re-worked the 𝐴𝐶 protocol. Previously, the mid-line main body of the 𝐴𝐶 was the seed with targets either side and stops at the amygdala. Now, we use a temporal pole as the seed, the main body of the 𝐴𝐶 as a waypoint and the contralateral temporal pole as the final target. For the human, we use the Harvard-Oxford temporal pole ROI (*56*). For the macaque, we use the CHARM temporal pole ROI (*54*). The temporal pole seed/target pair are flipped and tractography is repeated, taking the average of runs. Compared to the previous version, this protocol provides greater symmetry in resultant reconstructions and greater connectivity to the poles of the temporal cortex, as suggested in the literature (*96, 97, 98, 10*), and slightly enhanced connectivity to the amygdala.

##### Fornix (𝐹𝑋)

For the 𝐹𝑋, a main output tract of the hippocampus, we have added an exclusion mask to the amygdala to prevent 𝐹𝑋 leakage to the amygdala, thus providing a cleaner 𝐹𝑋 compared to the original XTRACT implementation (*31*). For the human protocol, we used the Harvard-Oxford amygdala ROI (*55*). For the macaque, we used the SARM amygdala ROI (*99*).

### 4.2 Data

#### 4.2.1 Macaque MRI data

We used six high quality ex-vivo rhesus macaque dMRI datasets, available from PRIME-DE (*100*). As described in (*35, 31*), these were acquired using a 7T Agilent DirectDrive console, with a 2D diffusion-weighted spin-echo protocol with single-line readout protocol with 16 volumes acquired at b=0 𝑠/𝑚𝑚^2^, 128 volumes acquired at b=4000 𝑠/𝑚𝑚^2^, and a 0.6 mm isotropic spatial resolution.

#### 4.2.2 Human MRI data

We used high quality minimally preprocessed (*101*) in-vivo dMRI data from the young adult Human Connectome Project (HCP) (*57, 28*). The HCP data were acquired using a bespoke 3T Connectom Skyra (Siemens, Erlangen) with a monopolar diffusion-weighted (Stejskal-Tanner) spin-echo echo planar imaging (EPI) sequence with an isotropic spatial resolution of 1.25 mm, three shells (b values = 1000, 2000, and 3000 𝑠/𝑚𝑚^2^), and 90 unique diffusion directions per shell plus 6 b = 0 𝑠/𝑚𝑚^2^ volumes, acquired twice with opposing phase encoding polarities. Data correspond to total scan time per subject of approximately 55 minutes. For this study, we randomly drew 50 HCP subjects (age range, 22-36 years of age, 24/26 females/males.

To assess robustness against data quality, we also used data from the UK Biobank (3T Prisma, 32 channel coil, 2 mm isotropic resolution, b values = 1000 and 2000 𝑠/𝑚𝑚^2^, 50 directions per shell). The UK Biobank data are acquired with approximately 6.5 minutes scan time per subject and therefore represent more standard quality datasets, achievable in a clinical scanner (*58*). Fifty subjects were randomly drawn from the UK Biobank (UKB) (age range 42-65 years of age, 31/19 females/males). For both HCP and UKB cohorts we ensured that the distribution of QC metrics (such as subject motion and image SNR/CNR) was representative of the full HCP and UKB cohorts that we had available.

#### 4.2.3 Macaque tracer data

Tracer data were used to test aspects of the striatal bundle protocols (Figure 3). These were made available by SRH and were obtained from an existing collection of injections in 19 macaque brains, from (*102, 103, 104, 105, 106, 62, 107, 48, 108, 50, 109*) and cases from the laboratory of SRH. Specifically, anterograde tracers were injected across 78 cortical locations and their terminations within the putamen were recorded in coronal slices of the NMT template space at 0.5 mm resolution. Specifically, the injection sites were first assigned to one of four cortical ROIs (frontal, parietal, temporal and sensorimotor cortices), obtained from the NMT CHARM v1 parcellation (*54*). For each of these four injection ROIs, we counted all the corresponding terminations within the putamen, and then divided by the total number of termination sites. This resulted in a termination probability map for each cortical region across the putamen, and these termination maps were smoothed using spline interpolation. The putamen mask was obtained from the NMT SARM v1 parcellation (*110*). To compare against tractography in F99 space, these maps were non-linearly registered from NMT to F99 space using RheMAP (*111*).

### 4.3 MRI data preprocessing

#### 4.3.1 Crossing fibre modelling and tractography

For both the human and macaque data, we modelled fibre orientations for up to three orientations per voxel using FSL’s BEDPOSTX (*112, 113*). These orientations were used in tractography. Probabilistic tractography was performed using FSL’s XTRACT (*31*), which uses FSL’s PROB-TRACKX (*114,115*). The standard space protocol masks were used to seed and guide tractography, which occurred in diffusion space for each dataset. 60 major white matter fibre bundles were reconstructed (30 cortico-cortical, 29 cortico-subcortical, 1 cerebellar, Supplementary Table S1). A curvature threshold of 80° was used, the maximum number of streamline steps was 2000, and subsidiary fibres were considered above a volume fraction threshold of 1%. A step size of 0.5 mm was used for the human brain, and a step size of 0.2 mm was used for the macaque brain. Resultant spatial path distributions were normalised by the total number of valid streamlines.

#### 4.3.2 Registration to standard space

For the human data, nonlinear transformations of T1-weighted (T1w) to MNI152 standard space were obtained. The distortion-corrected dMRI data were separately linearly aligned to the T1w space, and the concatenation of the diffusion-to-T1w and T1w-to-MNI transforms allowed diffusion-to-MNI warp fields to be obtained. For the macaque, nonlinear transformations to the macaque F99 standard space were estimated using FSL’s FNIRT (*116*) based on the corresponding FA maps.

For cases where NMT-space tractography protocols were used, nonlinear transformations to NMT space were obtained using RheMAP (*111*).

### 4.4 Tractography against data quality and individual variability

#### 4.4.1 Varying data quality

To explore robustness against varying data quality, we compared tractography reconstructions for in-vivo human dMRI data of considerably different data resolutions, diffusion contrast and scan time. Specifically, we explored whether tract reconstructions in state-of-the-art HCP data (approximately 55 minutes of scan time) were similar to reconstructions in bog standard data from the UK Biobank (UKB) (approximately 6.5 minutes of scan time), both on group average maps, as well as individual reconstructions.

Inter-subject variability for each tract reconstruction was assessed within and across the HCP and UKB cohorts. Inter-subject Pearson’s correlations were obtained by cross correlating random subject pairs tract-wise. Specifically, for each subject pair, we correlated the normalised path distributions in MNI space for each tract, after thresholding the path distribution at 0.5% (*31*), and then averaged the correlation across tracts for each subject pair. This was repeated for all possible unique subject pairs within and across cohorts.

A pairwise Mann-Whitney U-test was performed to determine differences in variability across analyses. For example we compared the HCP vs UKB correlation between original and the new(+revised) tracts. We corrected for multiple comparisons using Bonferroni correction.

We also explored tract reconstructions on individual subjects. To demonstrate representative results, we ranked subjects based on their tractography results against the cohort average and picked the the 10^𝑡ℎ^, 50^𝑡ℎ^ (median) and 90^𝑡ℎ^ percentile of the subjects. Specifically, for each subject, we calculated the average Pearson’s correlation value, to the group average, across all tracts. We then ranked the subjects based on this value.

#### 4.4.2 Respecting similarities stemming from twinship

As an indirect way to explore whether the proposed standardised protocols respected individual variability, we tested whether tractography reconstructions reflected similarities stemming from twinship. We used the family structure in the HCP cohort, to explore whether tracts of monozygotic twin pairs were more similar compared to tract similarity in dizygotic twins and non-twin sibling pairs, and to tract similarity in unrelated subject pairs, as would be expected by heritability of structural connections (*59,117,60*). We used the 72 pairs of monozygotic twins (MZ) available in the HCP cohort, and randomly selected 72 pairs of dizygotic twins (DZ), 72 pairs of non-twin siblings and 72 pairs of unrelated subjects, to have a balanced comparison. We compared tracts across pairs to assess whether our automated protocols respect the underlying tract variability across individuals. Specifically, for a given subject pair and a given tract, we calculated the Pearson’s correlation between the normalised path distributions (in MNI space and following thresholding at 0.5%). We repeated for all tracts and then calculated the mean correlation and standard deviation across tracts for that subject pair. This was then repeated for each group of subject pairs giving a distribution of average correlations for each group. We subsequently compared these distributions between the different groups. We repeated this process separately for the Original XTRACT tracts (*31*) and the new cortico-subcortical tracts to ensure that patterns were similar. For each analysis, a pairwise Mann-Whitney U-test was performed for all cohort pairs to determine the significant differences between them. We corrected for multiple comparisons using Bonferroni.

### 4.5 Building connectivity blueprints in cortex and subcortex

Connectivity blueprints are 𝐺𝑀 ×𝑇𝑟𝑎𝑐𝑡𝑠 matrices that have been proposed to represent the pattern of connections of GM areas to a predefined set of WM tracts (*35, 36*). To do so, the intersection of the core of WM tracts with the WM-GM boundary needs to be identified. For cortical GM, simply obtaining the intersection from the spatial path distribution maps of each tract would be dominated by the gyral bias in tractography near the cortex (*118*). Instead, whole-brain tractography matrices can be used as intermediaries. Specifically a 𝐺𝑀 × 𝑊 𝑀 connectivity matrix can be generated by seeding from each location of the white-grey matter boundary and targetting to a whole WM mask and this can then be multiplied by a 𝑊 𝑀 × 𝑇𝑟𝑎𝑐𝑡𝑠 obtained by collating the path distributions of all tracts of interest.

The cortical blueprints 𝐺𝑀_𝑐𝑡𝑥_ × 𝑇𝑟𝑎𝑐𝑡𝑠 were generated using our previously developed tool xtract blueprint (*35, 38*). We used the GM-WM boundary surface, extracted using the HCP pipelines (*101*) for the human data and the approach in (*35*) for the macaque. Briefly, a single set of macaque surfaces were derived using a set of high-quality structural data from one of the macaque subjects. The remaining macaque data were then nonlinearly transformed to this space, and the surfaces were nonlinearly transformed to the F99 standard space. All surface data were downsampled to 10,000 vertices prior to tractography. Volume space white matter targets were downsampled to 3 mm isotropic for the human and 2 mm isotropic for the macaque.

We extended the blueprint generation to include the subcortex. For subcortical nuclei we found that using an intermediary 𝐺𝑀 ×𝑊 𝑀 matrix did not help (as gyral bias is not relevant in subcortex - in fact it made patterns less specific). Hence, subcortical 𝐺𝑀_𝑠𝑢𝑏_ × 𝑇𝑟𝑎𝑐𝑡𝑠 blueprints were built using the intersection of the path distribution of each tract with the subcortical structures of interest (i.e. through multiplication of white matter tracts and binary subcortical masks, including putamen, caudate, thalamus, hippocampus, amygdala). Figure 9) shows a comparison of the two approaches for various tracts in the human and macaque: i) using an intermediary 𝐺𝑀 × 𝑊 𝑀 matrix to obtain subcortical connection patterns, as done in (*35*) for cortical regions and ii) using directly the tractography path distributions. The latter approach gave more focal and specific patterns and was used here for the subcortical regions. Tracts were downsampled (at 2 mm for human and 1 mm for macaque), thresholded at 0.1%, and multiplied by the subcortical nuclei masks, and then vectorised and stacked to create a 𝐺𝑀_𝑠𝑢𝑏_ × 𝑇𝑟𝑎𝑐𝑡𝑠 matrix. These were then row-wise concatenated (i.e vertically) with the cortical blueprints to generate CIFTI-style blueprints with approximately 10,000 cortical vertices and approximately 5,000 subcortical voxels (per left/right hemisphere). Finally, connectivity blueprints were row-wise sum-normalised. Following subjectwise construction of connectivity blueprints, we derived group-averaged blueprints for macaques and humans.

**Figure 9:**
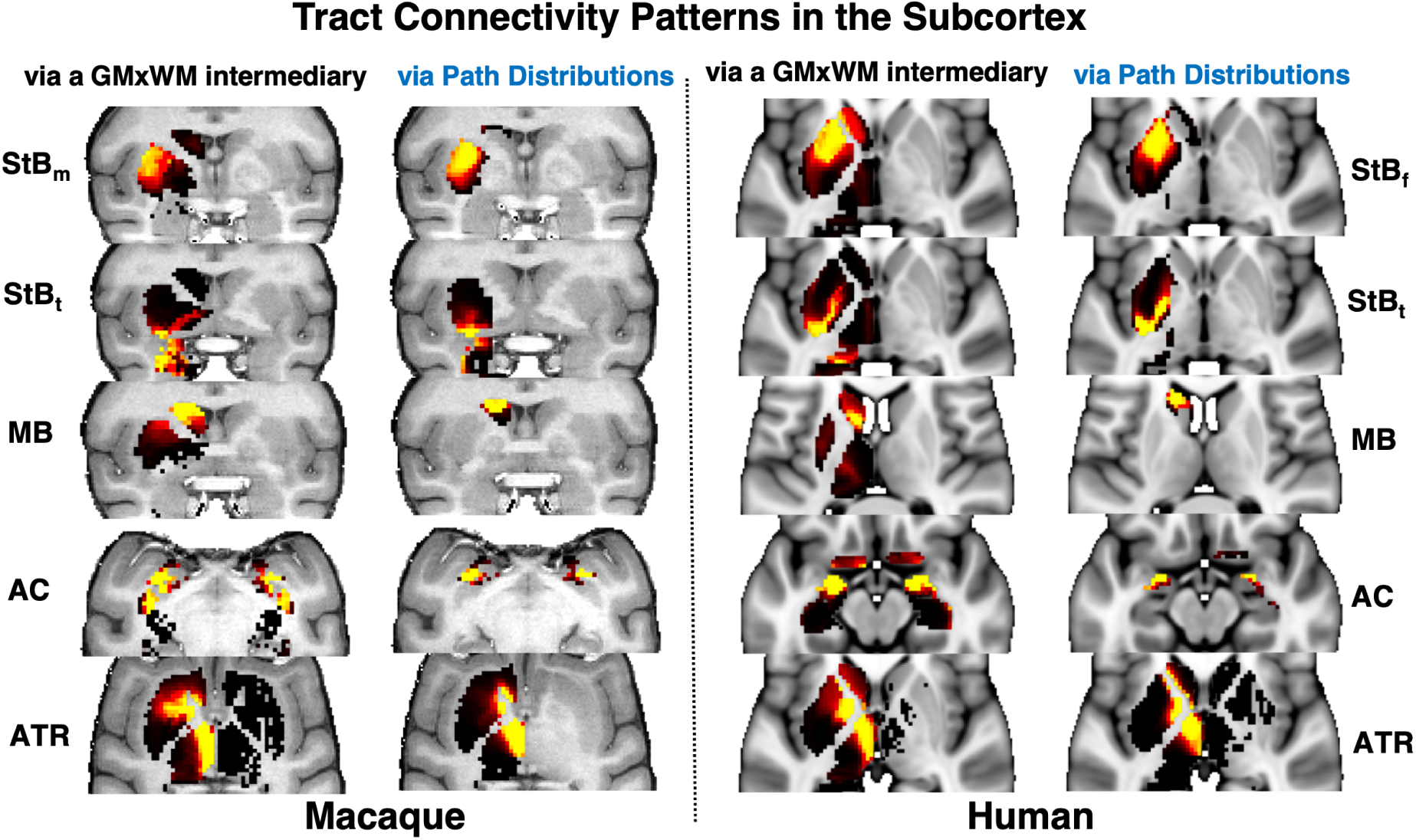
Improved specificity in subcortical connectivity patterns when using directly the tractography path distributions. Subcortical 𝐺𝑀_𝑠𝑢𝑏_ × 𝑇𝑟𝑎𝑐𝑡𝑠 blueprints were built using: i) an intermediary whole brain tractography 𝐺𝑀 × 𝑊 𝑀 matrix, multiplied by 𝑊 𝑀 × 𝑇𝑟𝑎𝑐𝑡𝑠 as done in (35) for cortical regions, ii) the intersection of the path distribution of each tract with the subcortical structures of interest. The two approaches are shown on the left and right columns for each of the macaque and human examples and for representative example tracts (rows). The latter approach resulted in improved specificity in both the macaque and human, with the tract of interest connecting more focally to the relevant subcortical nucleus. For instance 𝑆𝑡𝐵 tracts end up more specifically in the putamen, 𝑀𝐵 in the caudate, 𝐴𝐶 in the amygdala and 𝐴𝑇 𝑅 in the thalamus. All examples are shown as axial views, apart from 𝑆𝑡𝐵_𝑚_, 𝑆𝑡𝐵_𝑡_, 𝑀𝐵 in the macaque that are shown in coronal views.

### 4.6 Comparing connectivity blueprints across species

We compared grey matter connectivity patterns between humans and macaques (i.e. rows of the corresponding connectivity blueprint matrices), both in cortex and subcortex. As connectivity patterns are anchored by sets of homologously defined white matter landmarks, connectivity patterns may be compared statistically using Kullback-Leibler (KL) divergence (Equation 1) (*119*), as previously used (*35*).

Let 𝑀 be the macaque connectivity blueprint matrix, with 𝑀_𝑖𝑘_ linking grey matter (cortex or subcortex) location 𝑖 to tract 𝑘 = 1: 𝑇, with the set of tracts with length 𝑇. Let matrix 𝐻 be the equivalent matrix for the human brain. Vertices 𝑖 and 𝑗 in the macaque and human brains can then be compared in terms of their connectivity patterns 𝑀_𝑖𝑘_, 𝐻_𝑗 𝑘_, 𝑘 = 1: 𝑇 using the symmetric KL divergence 𝐷_𝑖 𝑗_ as a dissimilarity measure. To avoid degeneracies in KL divergence calculations induced by the presence of zeros, we shifted all blueprint values by 𝛿 = 10^−6^. We used the tool xtract divergence to perform all relevant calculations.

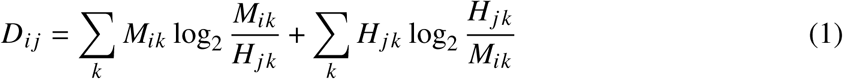

## Author contributions

All authors contributed significantly to this work from different and, in cases, overlapping roles. The specific contributions of each are: SA: Conceptualisation, Methodology, Data Curation, Formal Analysis, Investigation, Visualisation, Writing – Original Draft, Writing – Review and Editing SW: Methodology, Data Curation, Investigation, Writing – Review and Editing DF: Methodology, Formal Analysis, Writing - Review and Editing KB: Resources, Methodology, Writing - Review and Editing AMK: Resources, Methodology WT: Resources, Methodology, Writing - Review and Editing SJ: Methodology, Resources, Writing – Review and Editing SH: Resources, Methodology, Writing – Review and Editing RM: Methodology, Formal Analysis, Writing – Review and Editing SNS: Conceptualisation, Methodology, Formal Analysis, Resources, Writing – Original Draft, Writing – Review and Editing, Supervision, Funding Acquisition

## Data and materials availability

Human in-vivo diffusion MRI data are publicly available (www.humanconnectome.org/) and provided by the Human Connectome Project (HCP), WU-Minn Consortium (Principal Investigators: David Van Essen and Kamil Ugurbil; 1U54MH091657) funded by the 16 NIH Institutes and Centres that support the NIH Blueprint for Neuroscience Research; and by the McDonnell Centre for Systems Neuroscience at Washington University (*57*). The UK Biobank data were used under UK Biobank Project 43822 (PI: Sotiropoulos). Macaque data are openly available (https://fcon_ 1000.projects.nitrc.org/indi/PRIME/oxford2.html) and provided via the PRIMatE Data Exchange (http://fcon_1000.projects.nitrc.org/indi/PRIME/oxford2.html) (*100*).

Tractography protocols (https://github.com/SPMIC-UoN/xtract_data) and white matter tract atlases (https://github.com/SPMIC-UoN/XTRACT_atlases) will be made available on GitHub and will be released in FSL. Tools for performing standardised and automated tractography (XTRACT), building connectivity blueprints (xtract blueprint) and performing divergencebased comparisons of connectivity blueprints (xtract divergence) are available on GitHub (https://github.com/SPMIC-UoN/xtract) and are released in FSL (v6.0.7.10 onwards, https://fsl.fmrib.ox.ac.uk/fsl/docs/#/diffusion/xtract).

## Supplementary Materials

### Original cortico-thalamic XTRACT protocols

For completeness of presenting all the cortico-subcortical protocols, we include here the previously published protocols of thalamic radiations (*31*).

#### Acoustic Radiation (𝐴𝑅)

The acoustic radiation connects the medial geniculate nucleus (𝑀𝐺𝑁) of the thalamus to the auditory cortex. The seed was placed in the transverse temporal gyrus and the target covered the 𝑀𝐺𝑁 of the thalamus. The exclusion mask consisted of two coronal planes, anterior and posterior to the thalamus, and an axial plane superior to the thalamus. In addition, the exclusion mask contained the brainstem and a horizontal region covering the optic tract.

#### Anterior Thalamic Radiation (𝐴𝑇 𝑅)

The anterior thalamic radiation connects the thalamus to the frontal lobe. The seed mask was a coronal mask through the anterior part of the thalamus (*120*), with a coronal target mask at the anterior thalamic peduncle. In addition, the exclusion mask contained an axial plane covering the base of the midbrain, a coronal plane preventing leakage via the posterior thalamic peduncle and a coronal plane preventing leakage via the cingulum. A coronal stop mask covered the posterior part of the thalamus, extending from the base of the midbrain to the callosal sulcus.

#### Optic Radiation (𝑂𝑅)

The optic radiation consists of fibres from the lateral geniculate nucleus (𝐿𝐺𝑁) of the thalamus to the primary visual cortex. The seed was placed in the 𝐿𝐺𝑁 and the target mask consisted of a coronal plane through the anterior part of the calcarine fissure. Exclusion masks consisted of an axial block of the brainstem, a coronal block of fibres directly posterior to the 𝐿𝐺𝑁 to select fibres that curl around dorsally, and a coronal plane anterior to the seed to prevent leaking into longitudinal fibres.

#### Superior Thalamic Radiation (𝑆𝑇 𝑅)

The superior thalamic radiation connects the thalamus to the pre-/post-central gyrus respectively. The seed was a mask covering the whole thalamus and the target an axial plane covering the superior thalamic peduncle. An axial plane was used as a stop mask ventrally to the thalamus. The exclusion mask included two coronal planes, anterior and posterior to the target, to exclude tracking to the prefrontal cortex and occipital cortex respectively.

---

**Table S1:**
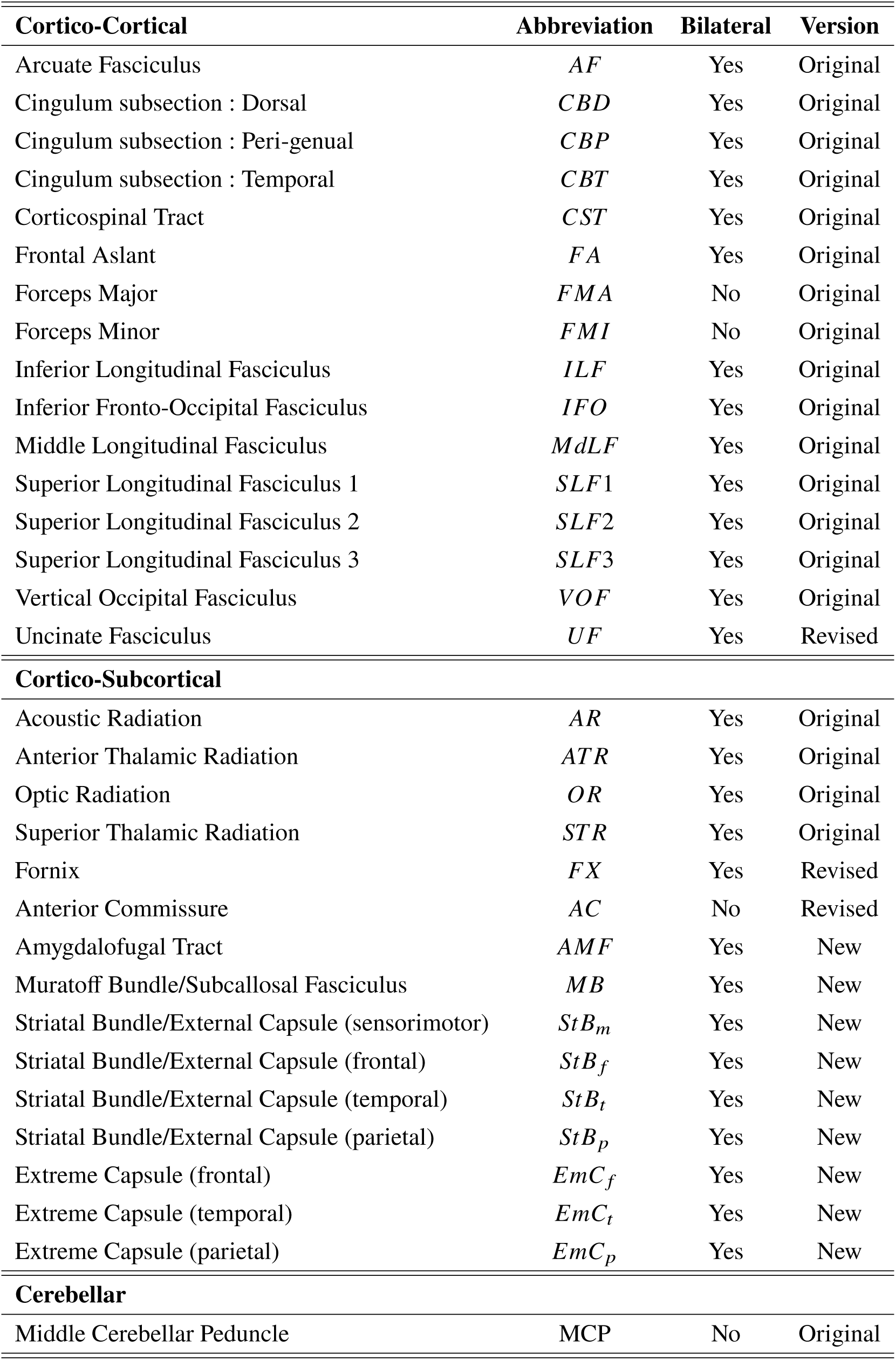
List of all species-matched (human & macaque) tract protocols. , grouped as cortico-cortical, cortico-subcortical and cerebellar. Columns indicate whether corresponding protocols are bilateral or not and whether they are New, Revised or not changed (Original XTRACT).

**Figure S1:**
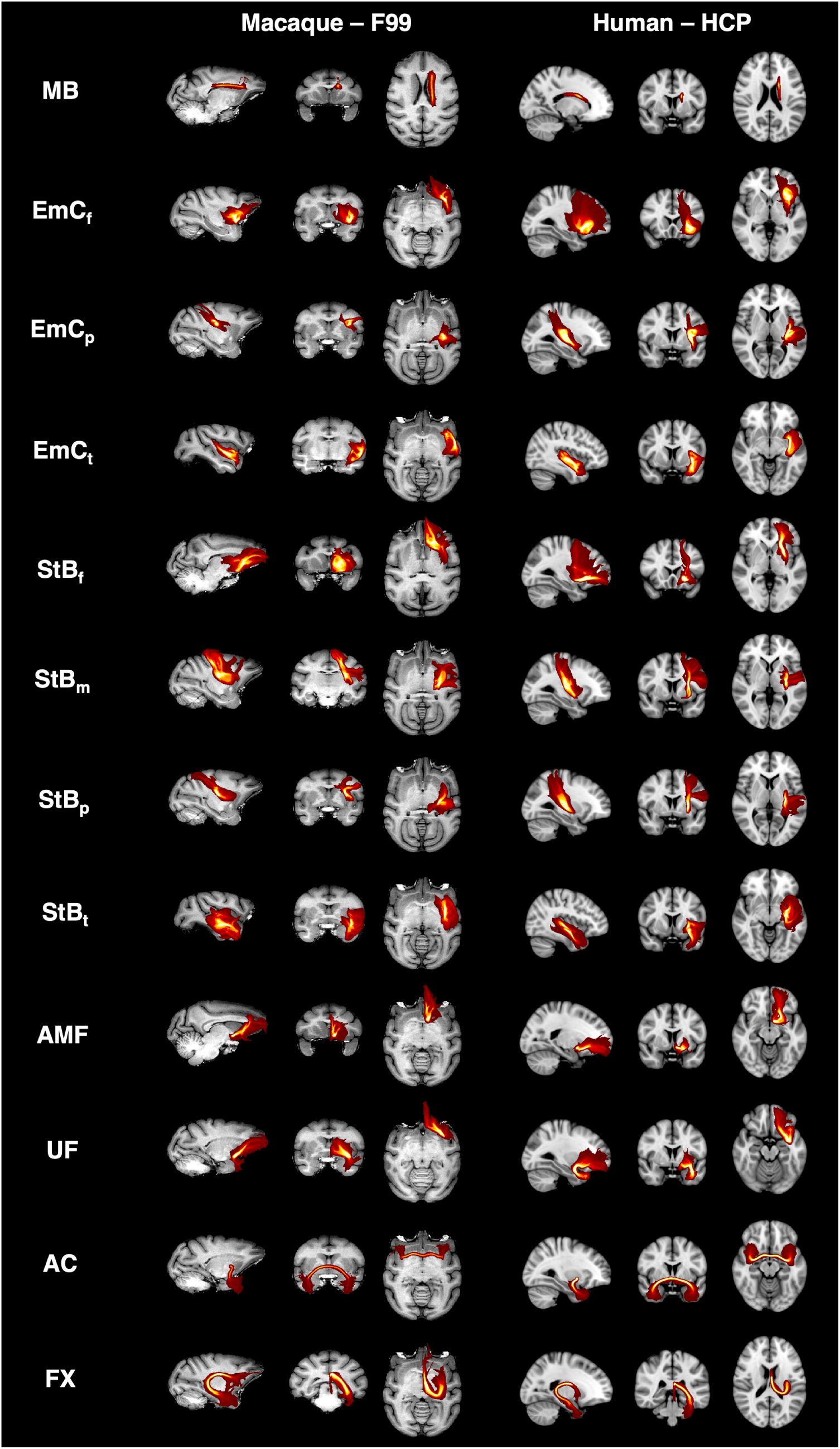
Tract reconstructions using our cortico-subcortical protocols, with good agreement between the macaque and the human. Maximum intensity projections (MIPs) of the group-averaged path distributions for all developed tractography protocols in the macaque (6 animal average) and human (50 healthy subject average from the Human Connectome Project dataset; HCP). All MIPs are across a window (20% of the field of view) centred at the displayed slices. Thresholded path distributions are displayed with a low threshold of 0.1% (for the 𝐸𝑚𝐶 parts the 90^𝑡ℎ^ percentile was used).

**Figure S2:**
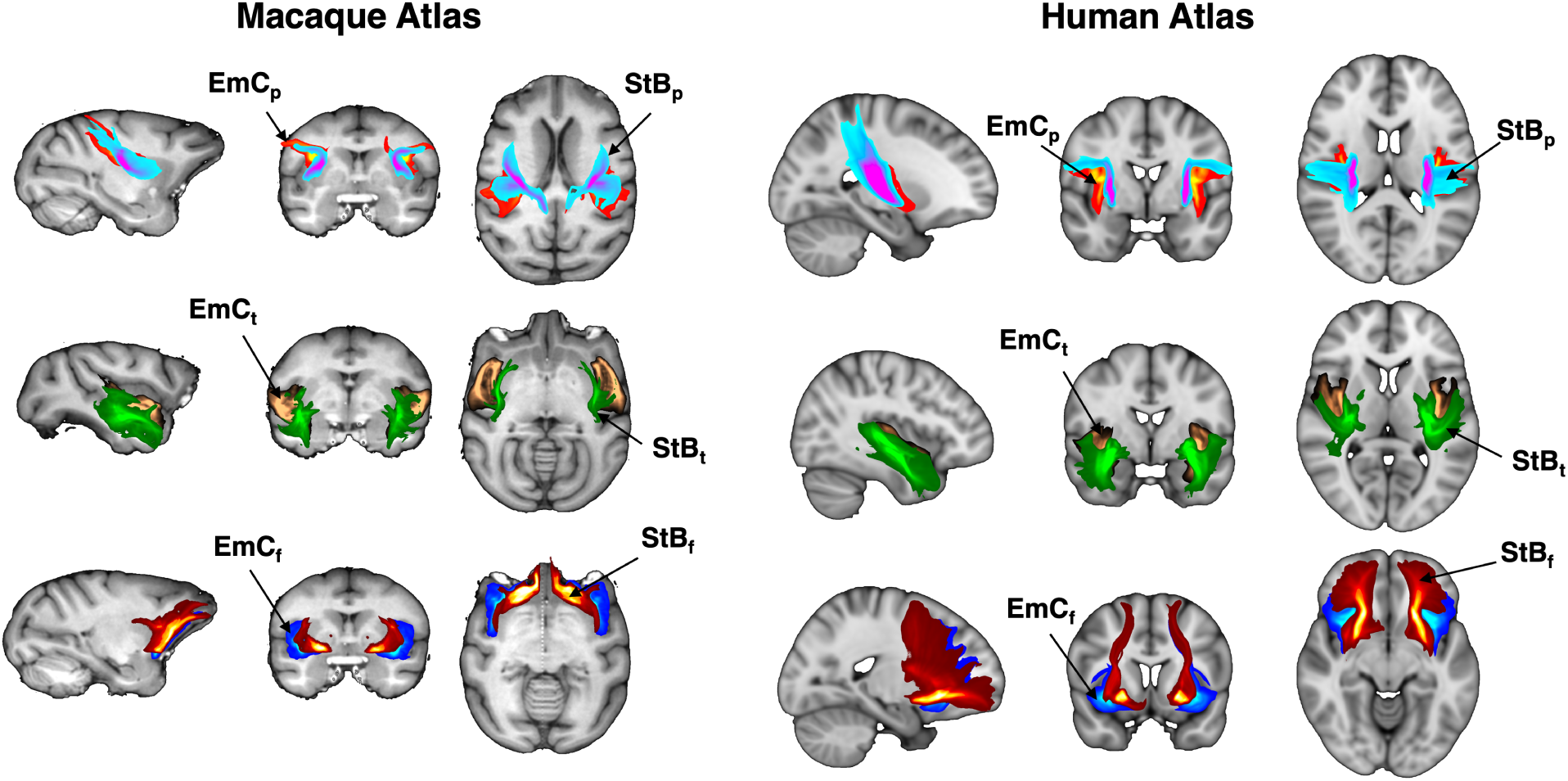
Relative positions maintained for all corresponding parts of striatal and extreme capsule bundles, across species. Maximum intensity projections (MIPs) of the group-averaged tractography results for corresponding parts of the striatal bundle (𝑆𝑡𝐵)/external capsule and the extreme capsule (𝐸𝑚𝐶) in the macaque (6 animal average) and human (50 healthy subject average from the Human Connectome Project dataset; HCP). For each part, 𝑆𝑡𝐵 is more medial and 𝐸𝑚𝐶 is more lateral (with respect to each other. Tracts considered: frontal, temporal and parietal parts of the anterior limb of the extreme capsule (𝐸𝑚𝐶_𝑓_, 𝐸𝑚𝐶_𝑡_, 𝐸𝑚𝐶_𝑝_*); frontal, temporal and parietal parts of the striatal bundle (*𝑆𝑡𝐵 _𝑓_, 𝑆𝑡𝐵_𝑡_, 𝑆𝑡𝐵_𝑝_*) (Table 1 in main text)*.

**Figure S3:**
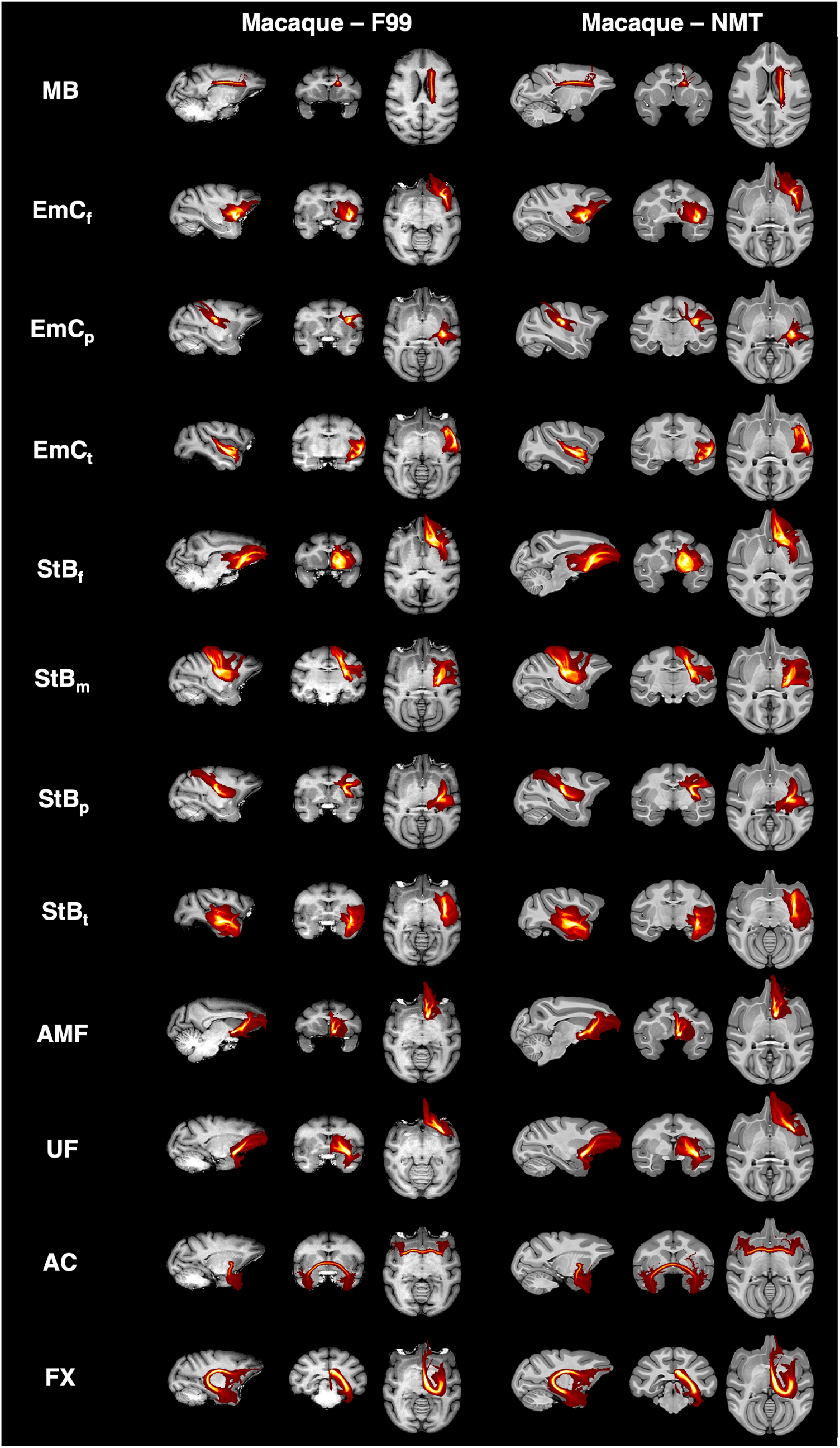
Corresponding tract reconstructions using our macaque cortico-subcortical protocols, between the two macaque standard spaces, F99 and NMT. Maximum intensity projections (MIPs) of the group-averaged path distributions for all developed tractography results for all developed protocols in the macaque (6 animal average) using protocols in the F99 standard space and protocols in the NMT standard space. All MIPs are across a window (20% of the field of view) centred at the displayed slices. Thresholded path distributions are displayed with a low threshold of 0.1% (for the 𝐸𝑚𝐶 parts the 90^𝑡ℎ^ percentile was used).

**Figure S4:**
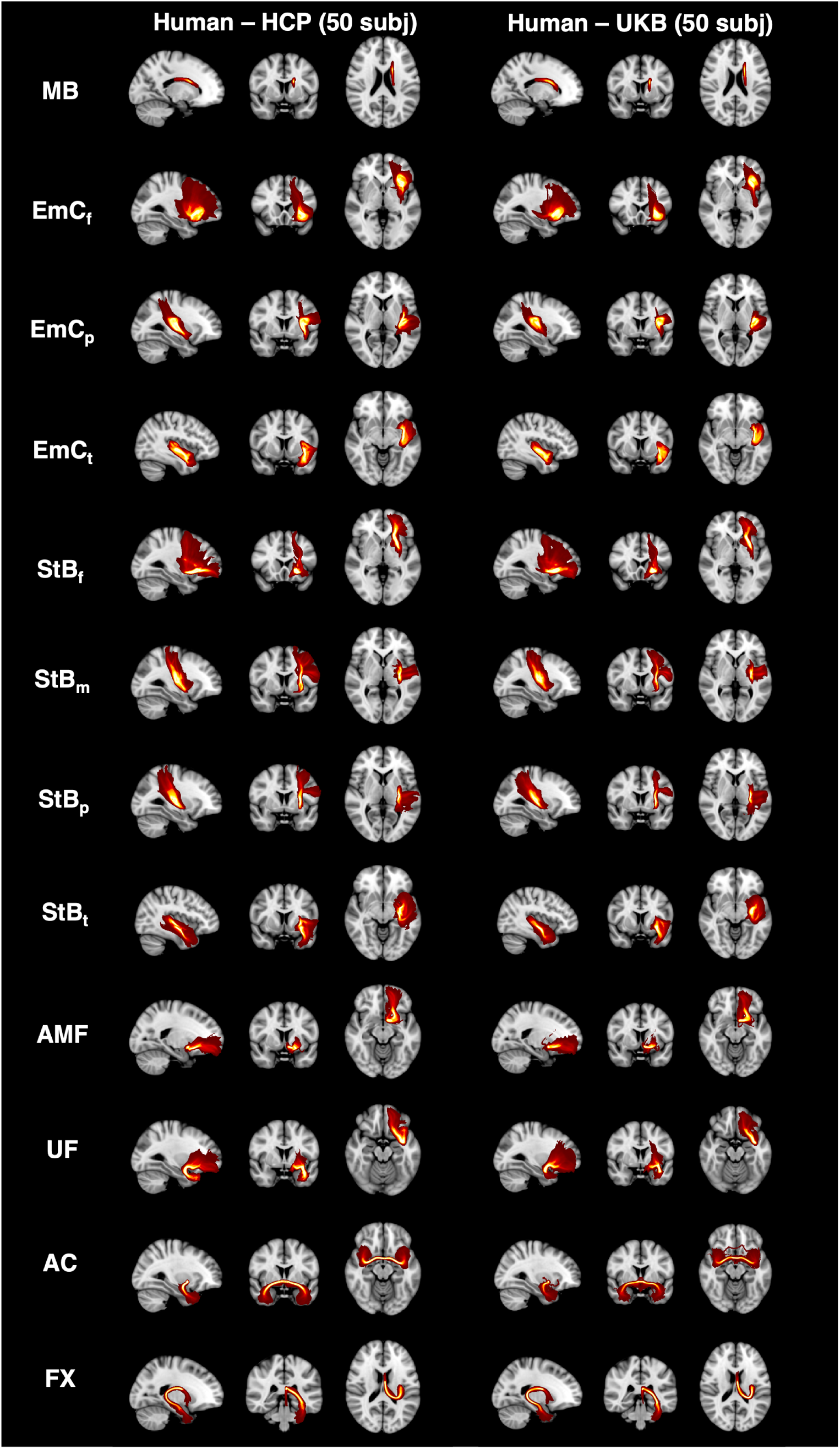
Corresponding tract reconstructions using our cortico-subcortical protocols, between different data resolutions (spatial and angular) and acquisition protocols. Maximum intensity projections (MIPs) of the group-averaged path distributions for all developed tractography results for all developed protocols in 50 Human Connectome Project (HCP) and 50 UK Biobank (UKB) subjects. All MIPs are across a window (20% of the field of view) centred at the displayed slices. Thresholded path distributions are displayed with a low threshold of 0.1% (for the 𝐸𝑚𝐶 parts the 90^𝑡ℎ^ percentile was used).

**Figure S5:**
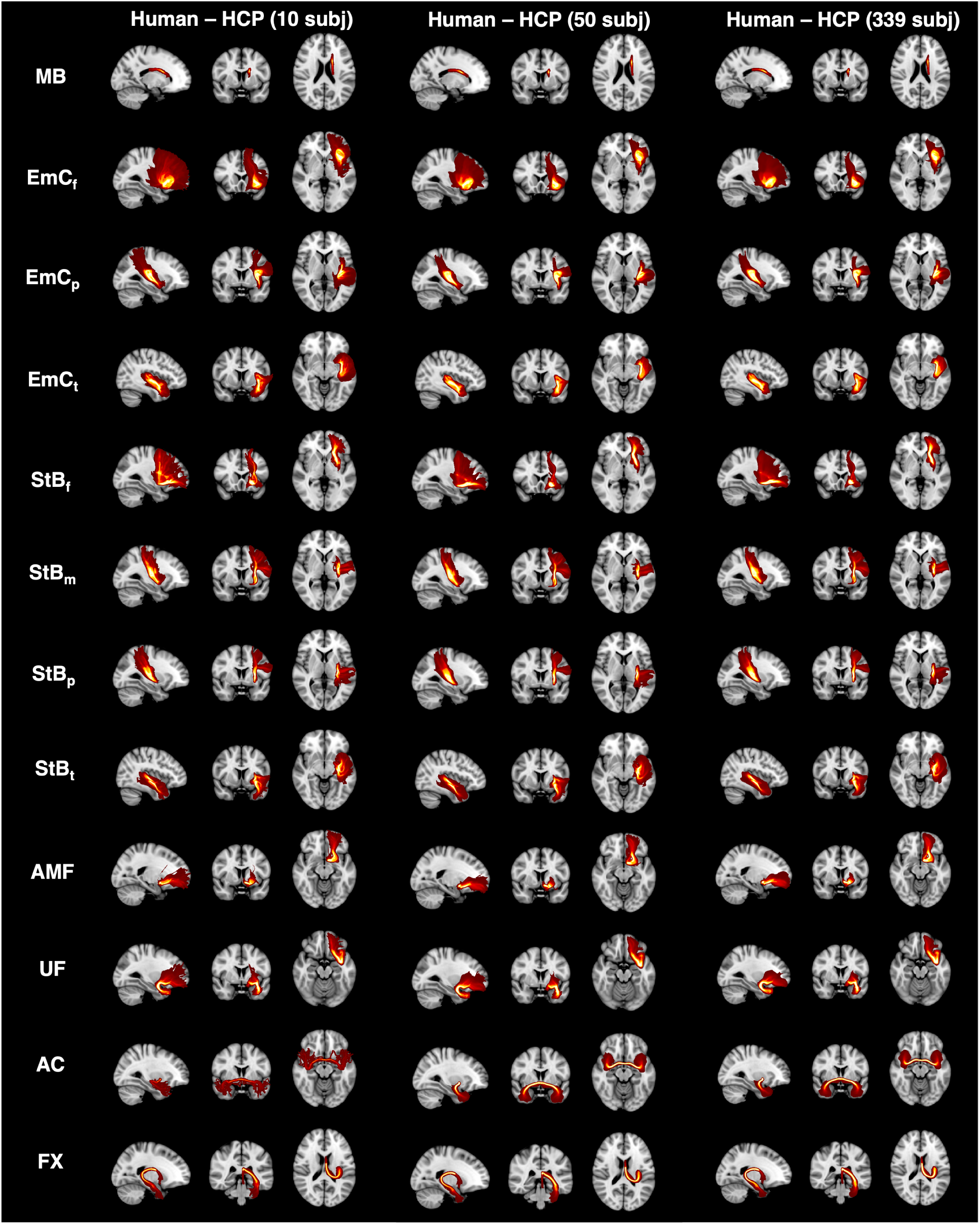
Corresponding tract reconstructions using our cortico-subcortical protocols, across different cohort sizes. Maximum intensity projections (MIPs) of the group-averaged path distributions for all developed tractography results for all developed protocols in 10, 50 and 339 (all unrelated subjects) Human Connectome Project (HCP) subjects. All MIPs are across a window (20% of the field of view) centred at the displayed slices. Thresholded path distributions are displayed with a low threshold of 0.1% (for the 𝐸𝑚𝐶 parts the 90^𝑡ℎ^ percentile was used).

**Figure S6:**
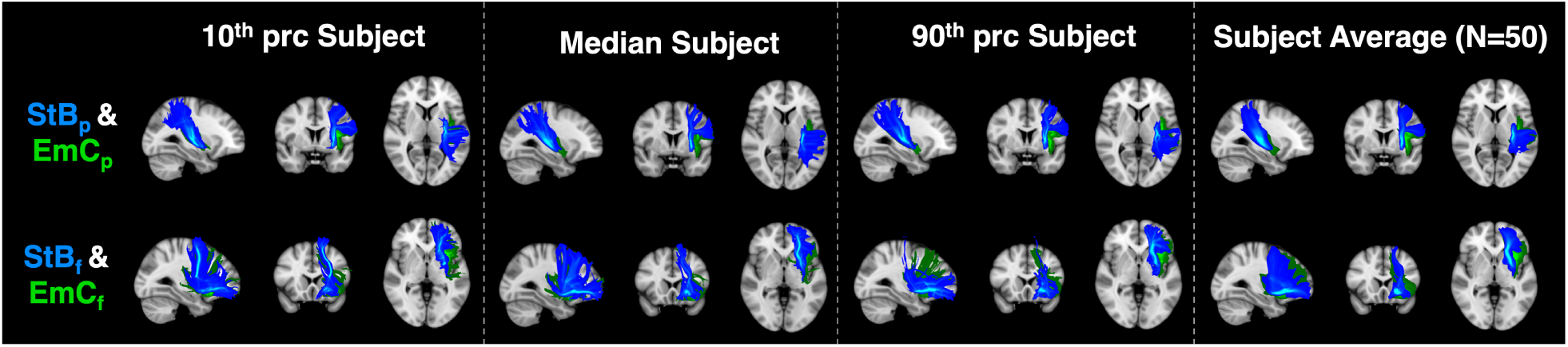
Individual subject tract reconstructions match the group average, whilst preserving expected topology for. 𝑆𝑡𝐵 **and** 𝐸𝑚𝐶. Showing reconstruction and relative topology for frontal and parietal parts (𝑆𝑡𝐵 _𝑓_ vs 𝐸𝑚𝐶_𝑓_, 𝑆𝑡𝐵_𝑝_ vs 𝐸𝑚𝐶_𝑝_) for individual subjects. The subjects chosen are those corresponding to the 10^𝑡ℎ^, 50^𝑡ℎ^ (median) and 90^𝑡ℎ^ percentile of the distribution of tract correlations against the group average for the HCP cohort (N=50). For each subject, the mean correlation to the average across all New tracts was computed and subjects were ranked based on this mean tract correlation. In all cases, tracts reconstruct similarly to the average atlas, whilst preserving the expected topology (𝑆𝑡𝐵 more medial than 𝐸𝑚𝐶).

**Figure S7:**
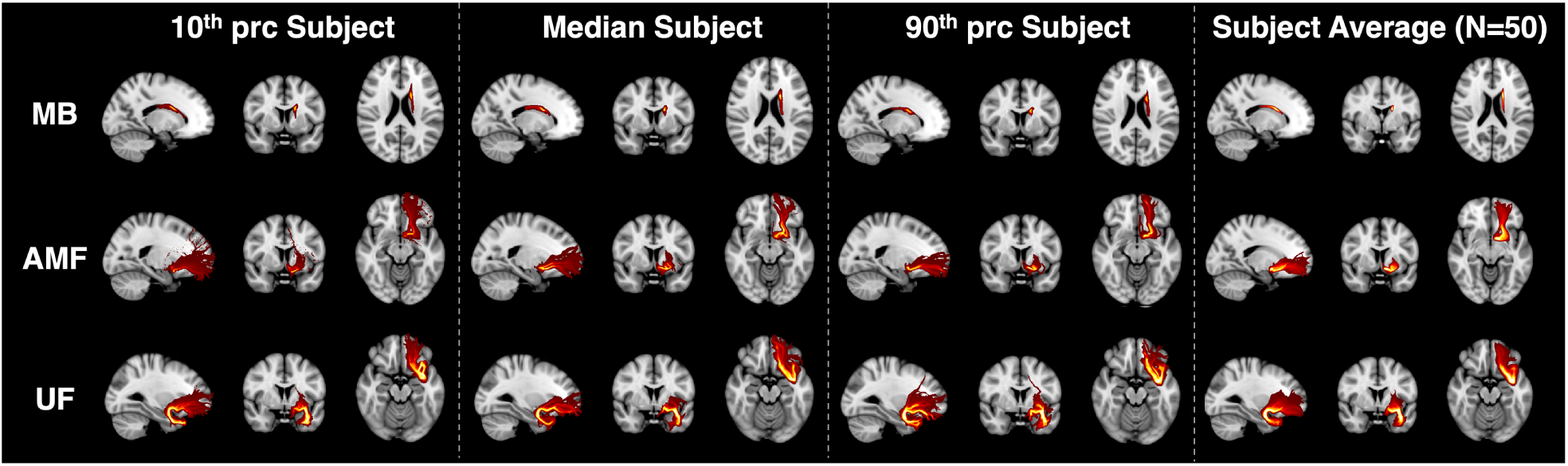
Individual subject tract reconstructions match the group average for. 𝑀𝐵, 𝐴𝑀𝐹 **and** 𝑈𝐹. The subjects chosen are those corresponding to the 10^𝑡ℎ^, 50^𝑡ℎ^ (median) and 90^𝑡ℎ^ percentile of the tract correlations against the group average for the HCP cohort (N=50). For each subject, the mean correlation to the average across all New tracts was computed and subjects were ranked based on this mean tract correlation.. In all cases tracts reconstruct similarly to the atlas.

